# Rebalancing Viral and Immune Damage versus Tissue Repair Prevents Death from Lethal Influenza Infection

**DOI:** 10.1101/2024.07.04.601620

**Authors:** Hiroshi Ichise, Emily Speranza, Federica La Russa, Tibor Z. Veres, Colin J. Chu, Anita Gola, Ronald N. Germain

## Abstract

Maintaining tissue function while eliminating infected cells is fundamental to host defense. Innate inflammatory damage contributes to lethal influenza and COVID-19, yet other than steroids, immunomodulatory drugs have modest effects. Among more than 50 immunomodulatory regimes tested in mouse lethal influenza infection, only the previously reported early depletion of neutrophils showed efficacy, suggesting that the infected host passes an early tipping point in which limiting innate immune damage alone cannot rescue physiological function. To re-balance the system late in infection, we investigated whether partial limitation of viral spread using oseltamivir (Tamiflu) together with enhancement of epithelial repair by blockade of interferon signaling or the limitation of further epithelial cell loss mediated by cytotoxic CD8^+^ T cells would prevent death. These treatments salvaged a large fraction of infected animals, providing new insight into the importance of repair processes and the timing of adaptive immune responses in survival of pulmonary infections.

## INTRODUCTION

Despite wide availability of antiviral drugs and vaccines, seasonal influenza can be life-threatening and lead to pneumonia and acute respiratory distress syndrome (ARDS), thus remaining a major public health threat that causes millions of hospitalizations and hundreds of thousands of deaths world-wide every year (*1*). Beyond this typical seasonal threat, influenza has caused multiple worldwide pandemics in the past 100 years (*2*) including the worst pandemic in recorded history, the 1918 H1N1 pandemic, which led to an estimated 50 million deaths (*3*).

Histopathology indicates that fatal influenza is associated with wide-spread pneumonia while modest cases are characterized by upper respiratory tract and tracheal infection, with diffuse alveolar damage, bronchitis, and bronchiolitis (*4*). Dysregulated innate immunity, leading to disruption of tissue homeostasis, is a major cause of the tissue damage in such infections (*5–8*). Pre-clinical studies have identified type I interferon (IFN) (*6*), neutrophils (*5, 9*), and CCR2^+^ monocyte-derived dendritic cells (*8*) as making major contributions to this damage. Excessive innate immune responses are also a hallmark of severe acute respiratory syndrome (SARS), Middle East respiratory syndrome (MERS), and coronavirus disease 2019 (COVID-19), highlighting further the relationship of poorly constrained innate immune responses to fatal pulmonary infection (*10–13*). Given these findings, anti-inflammatory treatments have been extensively studied in attempts to ameliorate severe influenza (*14*), but anti-viral treatment remains the only option recommended by the Centers for Disease Control and Prevention (*15*). For COVID-19, high-dose steroids have been found to be moderately effective (*16*), whereas again a wide range of immunosuppressive agents with greater selectivity have more modest effects at best when used late in the course of infection (*17*).

Temporal understanding of pathogenesis plays an increasingly important role in gaining the insights needed for rationale development of new clinical interventions in diverse disease states. A previous study showed that an early fatal gene signature associated with a neutrophil-mediated inflammatory feed forward circuit is seen at very early stages of influenza infection and acute depletion of neutrophils improves survival (*5*). Other research has shown that while type I interferons (IFN) play a major role in rapid anti-viral responses that limit pathogen spread, the timing of production of this cytokine is critical to preventing a pathogenic innate immune response in SARS-CoV infection (*18*). These findings raise the question of whether the limited success in delayed treatment of severe viral pneumonias using agents that modify the innate immune response can be traced to a timing issue – when such agents are used, it is possible that the damage caused by this inflammatory response may have already passed a tipping point that does not allow the return of a lung tissue physiological state solely through further limitation of damage from such pathways.

One of the earliest gene expression signatures accompanying the intense innate inflammatory response to severe influenza infection is that of tissue repair (*5*). This makes sense, as the host is attempting to compensate for the damage, seeking to maintain critical physiological function. We hypothesized that agents inhibiting innate inflammation, used late in the course of disease, may be ‘too little, too late’. However, it was also possible that the interventions already tested were simply not diverse enough to discover an effective agent. Here we tested over 50 regimes starting either early or in the mid-point of the disease course in a uniformly lethal model of influenza infection and again found that other than very early neutrophil depletion, no single immunomodulatory treatment had a measurable effect on survival. We therefore turned to a tipping point model in which, when treatment was instituted late in infection, the balance of repair and damage fell too far on the negative side to rescue the infected host. Type I IFNs can inhibit the alveolar type 2 (AT2) cell proliferation and differentiation (*19*), processes that are key features of the pulmonary epithelial repair process. It is also clear that late CD8^+^ T cell responses can further limit lung gas exchange capacity by killing infected alveolar type 1 (AT1) cells (*20*). For these reasons, we tested whether a combination of incomplete drug attenuation of viral spread plus either inhibition of type I interferon signaling or depletion of CD8^+^ T cells could rescue infected animals later in the disease course. In accord with these concepts and the tipping point model, late promotion of epithelial repair in concert with modest inhibition of viral replication led to significant increases in animal survival by rebalancing the net effect on pulmonary capacity. Likewise, eliminating the late damage to infected epithelial cells mediated by cytolytic CD8^+^ T cells also achieved salvage of infected animals. These findings show that strategies that enhance tissue repair while limiting late arising adaptive immune damage improve survival when treatment is delayed until late in the course of lethal influenza infections, providing a paradigm that should prove valuable in developing therapies for other pulmonary infections.

## RESULTS

### An oropharyngeal lethal influenza infection causes severe lung pathology

To develop a lethal influenza infection model that causes severe lung pathology, we utilized the oropharyngeal route and titrated the dose of influenza A/Puerto Rico/8/34 (PR8) used for infection (Figs. S1A and S1B), identifying the minimum viral load producing a uniformly lethal outcome. We settled on 100 TCID50 and undertook time-resolved, multiplex immunological profiling of the lungs, blood, bone marrow, and lung-draining lymph nodes of the infected animals (Fig. 1A). Following infection, mice showed ruffled fur, hunching, and strained breathing starting around 3 days post-infection (dpi). Viral RNA in the lung peaked at 2 dpi and animals displayed significant weight loss (p-value < 0.001) starting at 4 dpi, succumbing to infection between 8 and 10 dpi (Figs. 1B, S1C, and S1D).

**Fig. 1:**
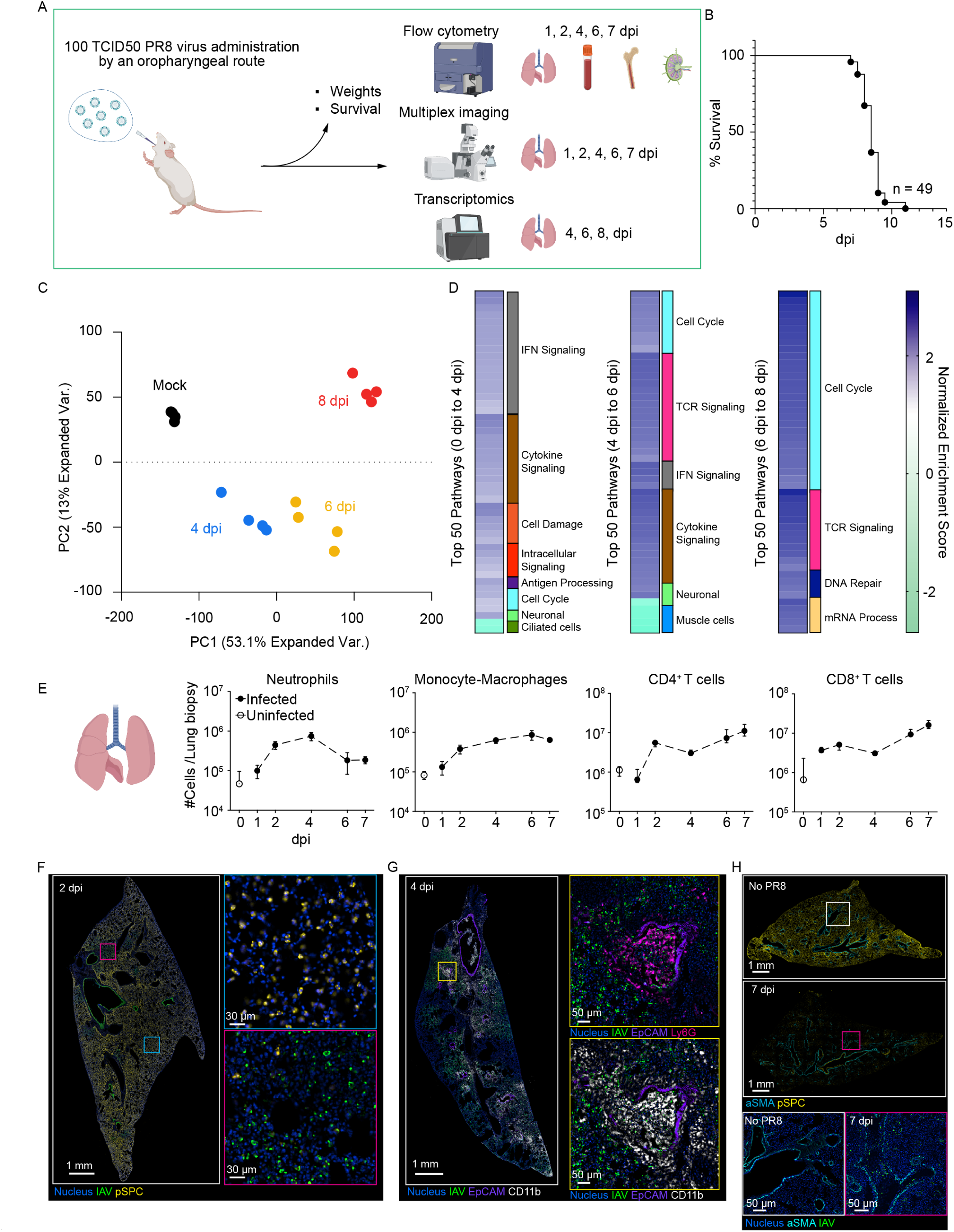
Characterization of lethal PR8 infection model. (A) Overview of the experimental setup. Mice were infected with 100 50% tissue culture infectious dose (TCID50) of PR8 via the oropharyngeal (OPh.) route, and body weight and survival were monitored. Infected animals were sacrificed on 1, 2, 4, 6, and 7 dpi and the lungs, peripheral blood, lung draining lymph node (LDLN), and bone marrow were collected for flow cytometry and imaging. Lung samples from 4, 6, and 8 dpi were used for RNA sequencing. (B) Kaplan-Meier plot of survival following PR8 infection. (C) PCA analysis of the global transcriptional changes from RNA-seq data at mock (black), 4 dpi (blue), 6 dpi(yellow), and 8 dpi (red) time points. (D) Groupings of the top 50 pathways based on adjusted p-value ranking from differential expression results at each day compared to the previous day. Values presented are the normalized enrichment score (NES) and are clustered into major groups based on the Jaccard index similarity between the genes in each pathway. (E) Flow cytometric analysis of the absolute number of different immune cell types in the lung samples. (E-G) Fluorescence microscopy imaging of lungs from control and infected animals at 2, 4, and 7 dpi. Colored squares show the region magnified in the panels to the right. Data are representative of at least 5 independent experiments with at least 7 animals in a group except for (C-E); data from one experiment.

To characterize the dynamics of the infection, RNA-seq was conducted on the whole lung at 4, 6, and 8 dpi. Reads associated with viral RNA matched with the qPCR results and showed an increase at 4 dpi that was sustained to 6 dpi. By 8 dpi, when animals are in a pre-terminal condition, the viral RNA began to decline (Fig. S1E). Transcriptional analysis of the time course showed early and strong signals associated with interferon signaling and inflammation by 4 dpi, followed by evidence of T cell responses at 6 dpi and a burst in cell proliferation at 8 dpi, suggesting ongoing repair processes (Fig. 1C and D). Additionally, cell type deconvolution of the transcriptional data showed early loss of AT2 cells by 4 dpi, with infiltration of neutrophils and macrophage-like populations at 6 dpi (Fig. S1F). These changes in cell type populations were confirmed by flow cytometric analysis showing sequential recruitment of neutrophil, myeloid, and T cell populations to the lung (Fig. 1E). Additional flow cytometric analysis highlighted the temporal differences in between lung and peripheral blood with respect to increases in these immune cell populations (Fig. S2).

Multicolor immunofluorescence microscopy confirmed the spread of the virus infection with increasing time after infection (Figs. 1F-1H, Fig. S3A). Importantly, infected AT2 cells exhibited diminished prosurfactant protein C (pSPC) expression that matched loss of *Sftpc* gene expression (FDR = 6 * 10^−8^) by 6 dpi (Fig. S3B). This correlated with the predicted decrease in AT2s by cell deconvolution at 4 dpi (adjust p-value < 0.0001), suggesting that tissue damage observed as a loss of pSPC presents from a very early phase in this PR8 infection model (Fig. 1F, S3C). Neutrophils and monocytes accumulated in the lungs and induced bronchiolitis at 4 dpi and persisted to 7 dpi (Fig. 1G). At 7 dpi, when animals are close to death, pSPC expression was completely lost, indicating severe pathology (Fig. 1H). Collectively, our model highlights distinct phases of virus- and immune cell-associated damage and severe pathology present in a fatal infection of PR8 influenza, including data consistent with the emphasis on innate inflammatory processes reported in prior studies.

### Early innate immunomodulation is not sufficient to rescue severe PR8 infection

Due to evidence in the literature and in the present knowledge that innate immune-associated damage may be a major factor contributing to influenza infection lethality (*5, 6, 8, 9*), we tested a wide spectrum of drugs that inhibit immune cell activity, signaling pathways, cytokines, and alarmins for their capacity to prevent death (Fig. 2A and table S1). We performed early or mid-course dosing of the drugs from 1-4 days post-infection with the goal of altering the immune response enough to overcome immunopathology causing lethality.

**Fig. 2:**
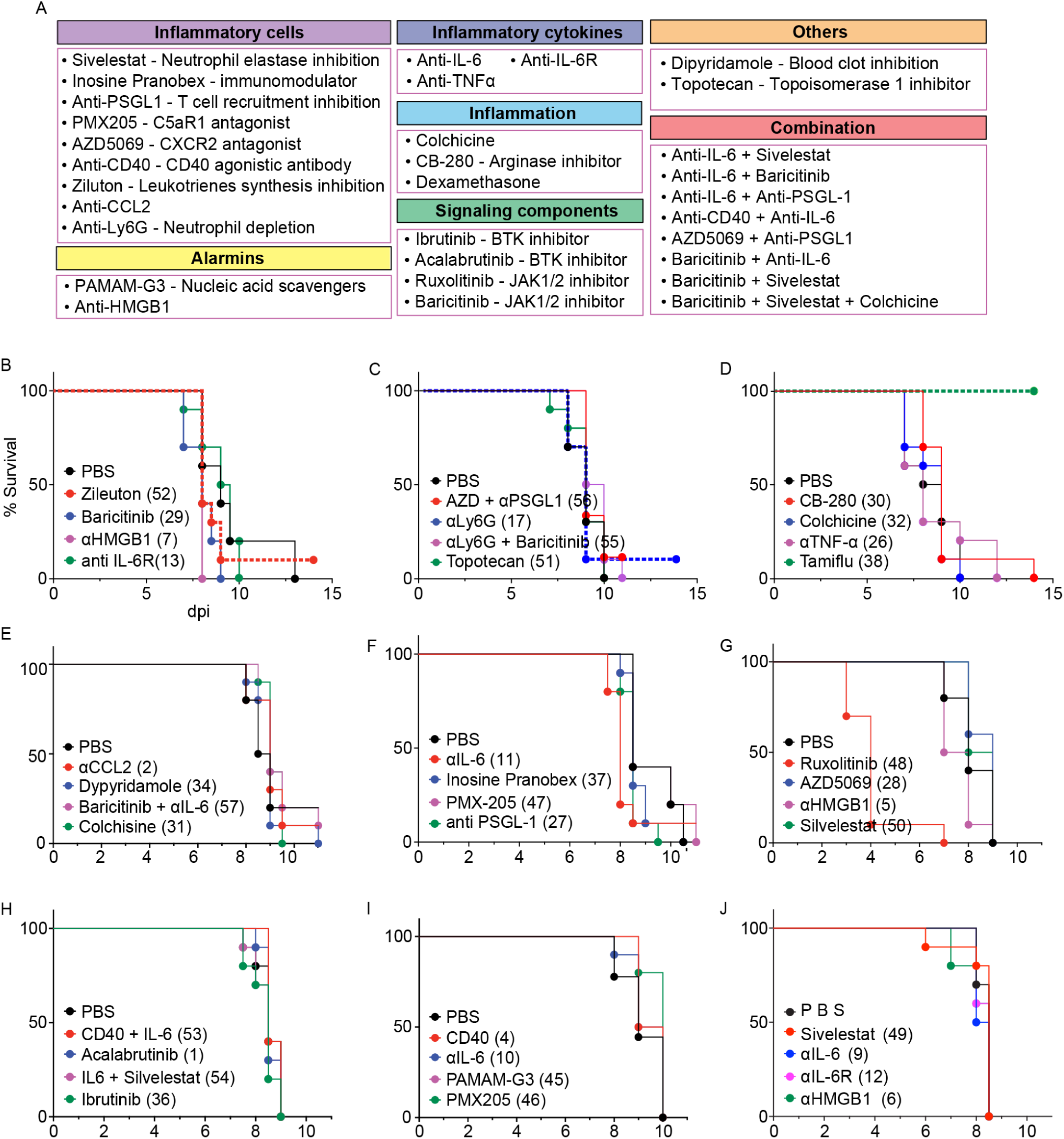
Early modulation of immunological/inflammatory pathways does not prevent death after infection. (A) Overview of the list of drugs used in this experiment. Drugs were administered as listed on table S1; survival was monitored. (B-J) Kaplan-Meier curves of groups treated with indicated inhibitors following 100 TCID50 PR8 infection. Numbers to the right of the drug name refer to the regimen listed in table S2. n=10 mice/group. Each graph represents an independent experiment with an independent PBS control.

Surprisingly, while we tested over 50 regimens (Fig. 2, table S2, and Fig. S4), only immunomodulatory treatments targeting neutrophils (Zileuton and anti-Ly6G depleting antibody) resulted in any survival (Fig. 2B and 2C), which was consistent with our previous findings (*5*). Treatment with an anti-viral drug (the neuraminidase inhibitor Tamiflu) (Figs. 2D) also allowed animals to survive infection. Inhibition of the virus resulted in complete survival of the treated cohort whereas inhibition of neutrophils resulted in minor increases in survival (10 % per group). While consistent with our previous findings of an early neutrophil-mediated inflammatory circuit as a cause of lethality-associated tissue damage in PR8 infection(*5*), the present findings show that interference with innate immune function is not sufficient to prevent death from infection even when used relatively early after infection or with combinations of drugs with different targets.

### Lethality-associated viral and immune damage occurs at very early stages

These drug screening results raised the important question of when and how damage leading to eventual death occurs and when and how this becomes apparently irreversible. In influenza infections, there are two different major pathways of tissue damage – immune-mediated tissue damage and direct cytopathic effect of the virus(*21*). To unravel the contributions of these two pathologic processes, we titrated anti-Ly6G antibody and Tamiflu to enable interference with innate immune-mediated(*5*) vs. virus-mediated tissue damage, respectively.

Titration of anti-Ly6G antibody showed a dose-dependent effect on host survival, though at higher concentrations of antibody than previously reported (Figs. S5A and S5B)(*5*). This form of antibody treatment had an incomplete effect on lung infiltration by neutrophils seen on day 3 dpi (Fig. S5C). However, more robust depletion of neutrophils with a combination of anti-Ly6G and anti-rat immunoglobulin κ light chain(*22*) as well as pre-infection depletion of neutrophils did not show a greater survival benefit, consistent with evidence that neutrophils not only cause damage but contribute to constraining viral damage (Figs. S5B and C). We then tried to dissect these two activities, seeking to determine the exact timing of when neutrophils were needed for survival vs. when they contributed to a damaging effect. To this end, we altered the timing of antibody treatment (Fig. 3A). Notably, compared to PBS treatment, neutrophil attenuation improved host survival only when administered on 1 dpi (p = 0.004) (Fig. 3B). Additionally, three-dimensional (3D) imaging of the lungs of untreated animals showed neutrophils filling some of the airways on 3 dpi (Movie S1). In line with the drop of total neutrophils in the lung at 3 dpi (Fig. S5C), histological analysis showed a significant improvement of bronchiole clogging (p-value = 0.016) (Figs. 3C and D) in animals treated with anti-Ly6G at 1 dpi. These results demonstrate that a cascade of immune-mediated damage caused by a neutrophil feed-forward circuit is completely irreversible at relatively early stages of infection and may also limit gas exchange via a large scale effect on bulk air movement in a manner similar to that reported for monocytes in SARS-CoV infection(*18*).

**Fig. 3:**
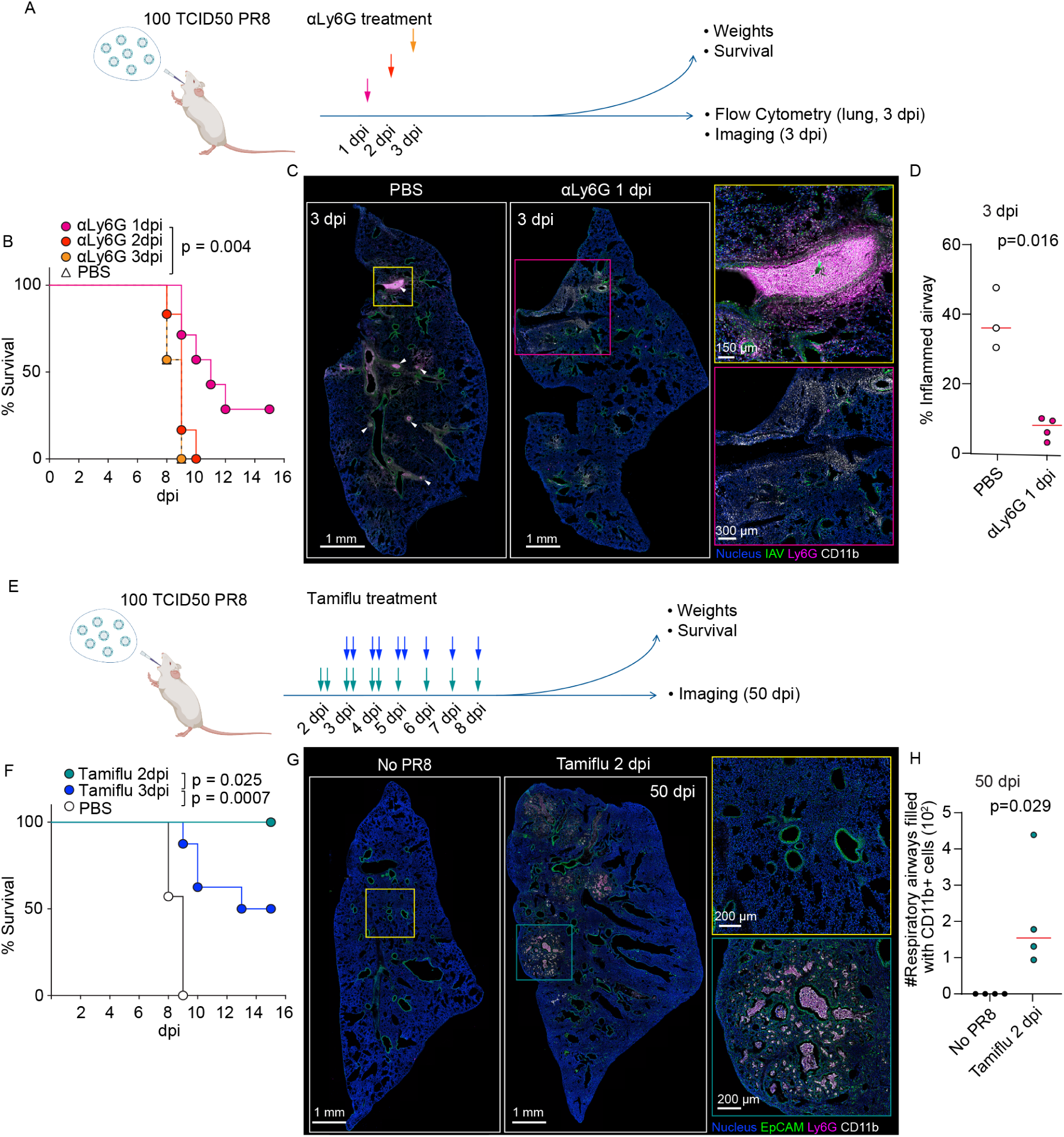
Irreversible viral and immune damage occur at very early stages of infection. (A) Overview of neutrophil depletion experiment. Mice were infected with 100 TCID50 of PR8 via the OPh. route and anti-Ly6G antibody was administered i.p. on either 1, 2, or 3 dpi. Weight and survival were monitored. (B) Kaplan-Meier curve of PR8 infected mice treated with anti-Ly6G antibody. 10 mice/group. (C) Flow cytometric analysis of neutrophil number in the lung biopsy on 3 dpi. Each dot represents an individual animal and the red bars represent the median. Data are representative of two independent experiments. (D) Representative immunofluorescence images of lungs from PR8-infected mice treated with anti-Ly6G antibody on 1 dpi. The lungs were harvested on 3 dpi. Magnified views of highlighted regions are shown on the right. (E) Overview of the Oseltamivir (Tamiflu) treatment experiment. Mice were infected as shown in (A) and Tamiflu was administered i.p. starting on either 2 or 3 dpi. Two doses per day were administered for the first 3 consecutive days with a single dose on the 4th and 5th day of administration. (F) Survival curve of PR8 infected mice treated with Tamiflu. Data are from one cohort with n = 7 mice/group. (G) Representative immunofluorescence images of lungs of PR8 infected mice treated with Tamiflu starting on 2 dpi. Magnified views of highlighted regions are shown on the right. The lungs were harvested on 50 dpi from animals surviving after Tamiflu treatment. (H) Quantification of myeloid infiltration in the alveoli shown in (G). Data are representative of two independent experiments unless specified.

After identifying when immune-mediated damage was occurring, we next addressed when the lethality-associated viral-mediated damage occurs by altering the day of Tamiflu administration (Fig. 3E). Starting Tamiflu treatment at 2 dpi, when the viral RNA reaches a peak, followed by daily administration of the drug, resulted in complete survival of animals (Fig. 3F). However, treatment starting on 3 dpi resulted in only partial survival, suggesting that existing viral-mediated damage at this early time in the context of intact immune responses to the infection has already reached an extent leading to eventual death despite inhibition of further viral spread. Strikingly, even in the group treated with Tamiflu starting at 2 dpi that resulted in 100% survival, irreversible lung damage is induced and leaves pathological sequelae detectable through 50 dpi, with scarring, loss of AT2 cells/cell function, and myeloid cell infiltration as dominant features (Figs. 3G and 3H). These results demonstrate that the cascade of lethality-associated tissue damage mediated by virus and the innate immune system are established at a very early phase of infection, before animals start losing body weight.

### Limiting immune or viral damage enhances tissue repair responses

Next, we addressed what biological processes are accelerated when we limit the viral- or immune-mediated damage early during infection. Since cell proliferation can be an indication of tissue repair(*23*) and untreated animals show evidence of a burst of proliferation at 8 dpi (Fig. 1C), we investigated if minimizing virus-associated tissue damage leads to earlier proliferation of epithelial cell layers. Consistent with the sequencing data, animals treated with PBS only showed a moderate increase in Ki-67 positive bronchial epithelial cells at 6 dpi compared to uninfected controls (adjusted p-value = 0.019). However, animals treated with Tamiflu on 2 dpi showed an increase in expression of Ki67 in the airway epithelium when compared to PBS treated animals at 6 dpi (adjusted p-value = 0.047) (Fig. 4A and 4B), indicating an increase of tissue repair when virus-mediated damage is limited. When animals were treated with anti-Ly6G antibody to limit neutrophil influx, there was a significant increase in Ly6C^+^Arginase-1^+^ macrophages in the lung at 6 dpi compared to PBS treated animals (p-value = 0.02) (Fig. 4C and 4D) in addition to reduced bronchiolitis (Fig. 3C and D). Arginase-1^+^ macrophages (known to be strong inducers of tissue repair(*24*)) were found in proximity to Ki67^+^ airway epithelial cells (Fig. 4C). These results demonstrate that ongoing viral propagation and immune responses can limit tissue repair.

**Fig. 4:**
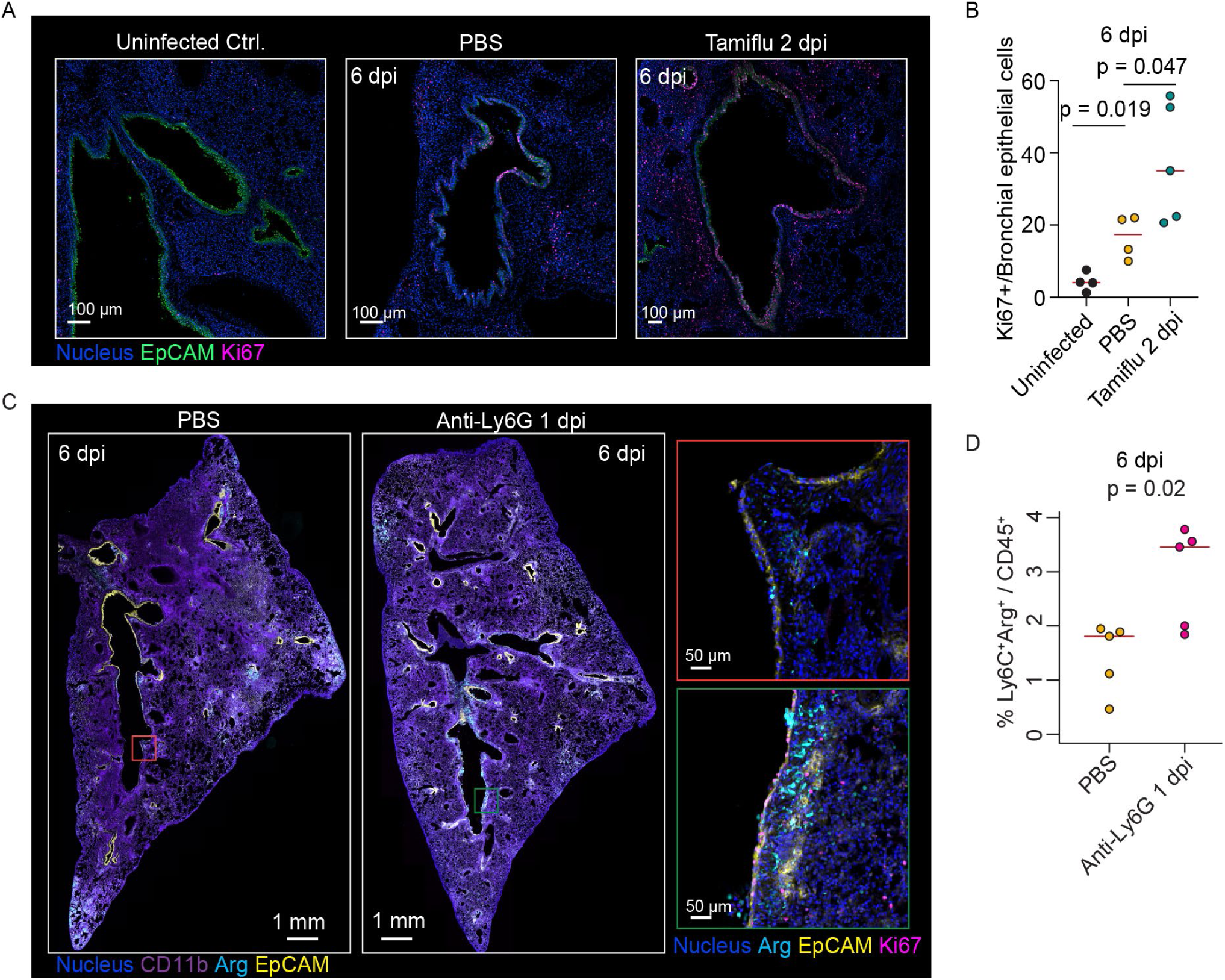
Limiting immune- or viral-mediated damage changes tissue repair responses. (A) Representative immunofluorescence images of lungs of PR8 infected mice treated with Tamiflu starting on 2 dpi. The lungs were harvested on 6 dpi. (B) Quantification of Ki67^+^EpCAM^high^ bronchial epithelial cells as shown in (A). (C) Representative immunofluorescence images of lungs of PR8 infected mice treated with anti-Ly6G antibody on 1 dpi. The lungs were harvested on 6 dpi. Magnified views of highlighted regions are shown on the right. (D) Quantification of Ly6C^+^Arginase-1^+^ cells in the lung on 6 dpi by flow cytometry. Each dot represents an individual animal and the red bars represent the median. Data are representative of two independent experiments.

### Combination treatments to enhance repair and limit both viral and late adaptive immune damage improve host survival

The preceding data suggested that the combination of viral and immune processes create sufficient damage at an early stage of infection such that the competing process of tissue repair is unable to maintain an adequate level of pulmonary function, even in the face of innate immune inhibitors that limit further damage. This led us to a ‘tipping point’ model in which survival past this damage checkpoint would require either enhancing repair to overcome the existing damage or limiting further pathology mediated by later arising adaptive, not innate, immune mechanisms such as cytotoxic CD8^+^ T cells, both under an umbrella of adequate viral inhibition. Previous studies revealed that type I IFN signaling pathways suppress proliferation and induce the death of AT2 cells during recovery from influenza infection (*19*). AT2 cells act as a reservoir of stem cells for replacement of AT1 cells in the presence of insults causing pulmonary epithelial damage (*25*). Furthermore, a post-mortem study of lung tissue from COVID-19 patients showed that the top signature obtained in a scRNAseq analysis was of stressed AT2 cells (*26*). In our model of lethal influenza, there is early and sustained expression of many type I IFN responsive genes (Fig. S6A), which may limit AT2 cell-dependent epithelial repair. At the same time, CD8^+^ T cells kill virus-infected cells, and this adaptive immune response late in infection could further deplete AT1 and AT2 cells that are crucial for gas exchange and epithelial repair (*25, 27–29*). Our evidence of T cell-associated signaling and accumulation of CD8^+^ T cells in the lung beginning at 6 dpi is correlated with further loss of surfactant proteins and the corresponding gene transcripts afterwards (Figs. 1D, 1E, S2B, and S2C), suggesting a negative impact of this adaptive immune response on lung function that can override ongoing repair. These findings led us to hypothesize that a counter-intuitive approach combining late inhibition of viral spread together with blocking of type I IFN signaling or depletion of CD8^+^ T cells could rebalance the damage-repair equation by promoting more efficient alveolar epithelial cell generation or preventing adaptive immune damage to the epithelial cell population. To test this hypothesis, we altered the timing of Tamiflu with clinically relevant doses to determine when Tamiflu alone could no longer effectively induce robust host survival. Unlike the previous high dose of Tamiflu (60 mg/kg), a clinically relevant dose (20 mg/kg) showed a more moderate effect even when used beginning on 2 dpi (Fig. S6B). Treatment starting on 4 dpi and 5 dpi showed comparable results (Fig. S6B) with around 40% survival and a delayed time to death (p-value = 0.003 and p-value = 0.002). Based on these data, we decided to proceed with the treatment starting with 4 dpi at 20 mg/kg, a protocol that mimics the clinically relevant scenario in which patients arrive to the hospital or seek treatment several days to a week after symptoms begin.

We simultaneously treated animals with either anti-IFNAR antibody or anti-CD8 antibody in addition to Tamiflu, treated with Tamiflu alone, or treated with PBS alone (Fig. 5A). A treatment effect was verified through RNA sequencing that monitored expression of type I IFN-induced genes or CD8a expression (Figs. S6D and S6E). In comparison to the dozens of agents targeted at the innate immune response that had failed to promote survival, both combination treatments significantly improved the survival rate compared to Tamiflu alone (p-value = 0.026 and p-value < 0.001) (Fig. 5B). Only the combination of Tamiflu with anti-IFNAR antibody showed improved body weight beginning at 12 dpi compared to Tamiflu alone (p-value 0.036) and this difference was maintained to 14 dpi (p-value = 0.037) (Fig. S6C). Notably, the combination of anti-IFNAR and anti-CD8 antibodies in addition to Tamiflu did not show a synergistic effect, but instead worsened outcome (Fig. 5B). This result indicates that either type I IFN signaling or a CD8^+^ T cell response is necessary for host defense, even in animals treated with Tamiflu. Neither individual treatment with anti-IFNAR or anti-CD8 antibodies showed a survival benefit (PBS vs anti-IFNAR, p-value = 0.94; PBS vs anti-CD8, p-value = 0.11), highlighting the necessity of limiting ongoing viral-mediated damage while treating to rebalance immune damage and epithelial repair.

**Fig. 5:**
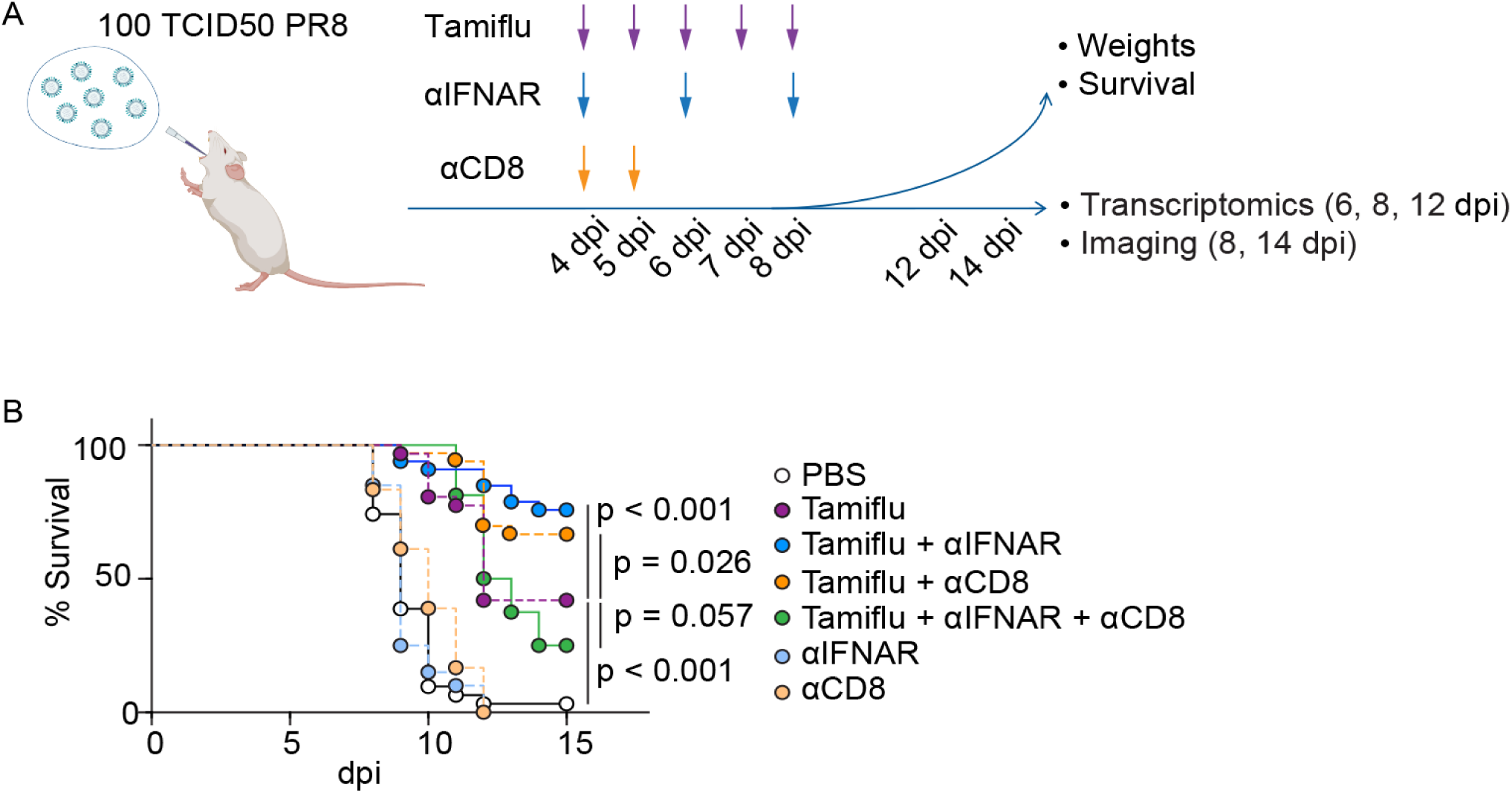
Combination strategies that target type-I IFN signaling or CD8^+^ T cell response in addition to viral-mediated damage improve host survival. (A) Overview of combination treatments to enhance repair and limit both viral and adaptive immune damage. Mice were infected with 100 TCID50 of PR8 via the OPh. route and Tamiflu and/or anti-CD8 antibody, and/or anti-IFNAR1 antibody were administered i.p. at the indicated time points. Weight and survival were monitored. RNA sequencing and immunofluorescence imaging were carried out at 4, 6, and 8 dpi and 8 and 14 dpi, respectively. (B) Kaplan-Meier plot of infected mice given PBS (n=32), Tamiflu (n=31), Tami + αIFNAR1 (n=33), Tami + αCD8 (n=33), Tami + αIFNAR1 + αCD8 (n=16), αIFNAR1 (n=20), and αCD8 (n=20). Log-rank test was used to calculate p values. Data are aggregated from 2-4 independent experiments.

### Blocking type I interferon signaling promotes early progenitor cell accumulation and enhanced epithelial repair

To investigate the mechanistic basis for the pro-survival effects of Tamiflu + anti-IFNAR or Tamiflu + anti-CD8, we carried out RNA-sequencing on whole lung tissues (Fig. 5A). Global principal component analysis (PCA) revealed that at 6 dpi the samples had similar transcriptional profiles. However, by 8 dpi, the Tamiflu + aIFNAR samples fell at a different location along principal component (PC) 2 (Figs. 6A, 6B, S7A, and S7B). This principal component was enriched for genes associated with cell proliferation, DNA repair, and cilia formation, consistent with enhanced repair processes in this group (Fig. S7C). Pathway analysis of all the conditions at 8 dpi compared to PBS treated animals showed expected patterns of downregulated type I IFN signaling in Tamiflu + anti-IFNAR samples and decreased T-cell receptor-associated signaling in Tamiflu + anti-CD8 samples (Figs. 6C and S8A). Across the many conditions, only 4 pathways were significantly up-regulated in any condition compared to PBS controls. In the Tamiflu + aIFNAR treated animals, pathways associated with keratinization were significantly enriched (adjusted p-value = 0.00003 and 0.001) (Fig. 6C). Assessment of genes associated with this pathway revealed three genes that were uniquely expressed in Tami + anti-IFNAR samples compared to PBS: *Krt14* (FDR = 0.003), *Trp63* (FDR = 0.04), and *Krt5* (FDR = 0.02) (Fig. 6D). These genes are known markers of basal epithelial cells during differentiation and show increased expression in the lung following injury, while keratin (KRT) 14^+^ p63^+^ KRT5^+^ cells can act as progenitors to damaged epithelial layers (*30*). In addition to this unique gene signature, the Tamiflu + anti-IFNAR condition showed a significant predicted enrichment of mesenchymal-like cells at 8 dpi compared to uninfected controls (adjusted p-value = 0.0487) (Fig. S8B). These cells play an important role in promoting proliferation of AT2 cells and regenerating the epithelial layers of the lung (*31, 32*). By 12 dpi, the Tamiflu alone samples had up-regulated similar gene programs, suggesting a delayed repair response due to ongoing interferon responses in the lungs. Together this suggests that the addition of anti-IFNAR antibody to Tamiflu therapy induces an earlier / enhanced repair program in the lungs and moves pulmonary function back into a physiologically supportive range.

**Fig. 6.**
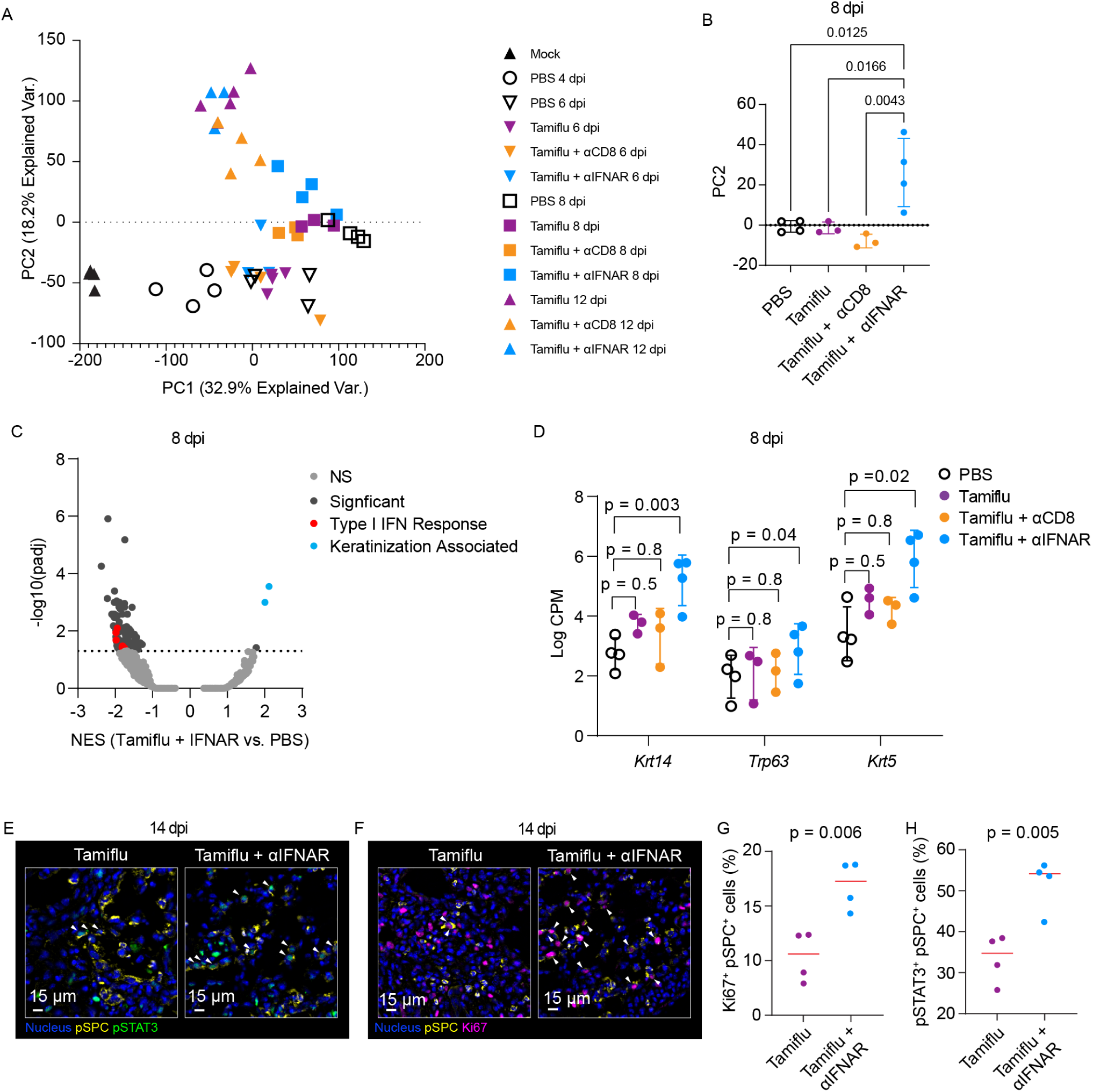
Combination strategies that target type-I IFN signaling in addition to viral-mediated damage enhance early tissue repair. (A) Principal component analysis of all samples and all conditions, showing the first two principal components. (B) Location of samples from 8 dpi along principal component (PC) 2. An ordinary one-way ANOVA was used to compare samples to Tamiflu + aIFNAR. (C) Pathway analysis of Tamiflu + aIFNAR compared to PBS only. The NES score is on the x-axis and the −log10(adjusted p-value) is on the y-axis. Pathway groupings are colored dots (NS = not significant). (D) Differential expression analysis of three genes in each condition compared to PBS. FDR are shown from the edgeR glmLRT test. (E – H) Immunofluorescence images and quantification of pSTAT3^+^ (E and G) or Ki67^+^ (F and H) AT2 cells in lungs from PR8 infected mice treated with Tamiflu or Tamiflu + anti-IFNAR1 antibody starting on 4 dpi. The lungs were harvested on 14 dpi. White arrows denote pSTAT3^+^pSPC^+^ cells in (E) or Ki67^+^pSPC^+^ cells in (F). Each dot represents an individual animal, and the red bars represent the median. Data are representative of two independent experiments except for (A-D); data from one experiment. n = 3-4 mice/group.

Multiplex imaging of tissues from animals given anti-IFNAR antibody with Tamiflu revealed significant maintenance of the bronchial epithelial cells at 8 dpi compared to controls treated with Tamiflu (p-value = 0.001) (Figs. S8C and S8D), consistent with the pro-repair basal epithelial cell differentiation gene signature. We next looked directly for evidence of proliferation and regeneration of AT2 cells at 14 dpi. STAT3 signaling in AT2 cells plays an important role in promoting alveolar epithelial regeneration via the BNDF-TrkB axis (*31*). The combination of anti-IFNAR antibody with Tamiflu significantly increased the phosphorylation of STAT3 as well as the expression of Ki67 in the AT2 population (Figs. 6E-6H). In line with these results, the absolute number of total and proliferating pSTAT3^+^ AT2 cells was higher in the animals treated with the combination of anti-IFNAR antibody and Tamiflu (Figs. S8E and F), showing that this treatment promotes an acceleration of alveolar epithelial cell regeneration crucial for improving pulmonary function. Together, these data show that inhibition of the type I IFN response promotes early stem-like cell accumulation that can induce repair of the epithelial layers and facilitate proliferation of AT2 cells, thus countering the effects of immune and virus-mediated damage.

### Inhibition of CD8^+^ T cell responses limits severe loss of AT2 cells

We next turned to an analysis of the effects of the anti-CD8 antibody treatment that together with Tamiflu was also successful in salvaging infected animals. In PBS treated animals, we see T cell responses starting to become enriched by 6 dpi (Fig. 1D). This is associated with a loss of surfactant gene expression (*Sftpc* p-value = 0.0067) (Fig. 7A) and loss of pSPC staining (Fig. 1H). When analyzing the expression of surfactant genes from the RNA-seq data across the time course, we observed an increased drop in expression at 8 dpi in both the Tamiflu alone treated animals and even more so in the Tamiflu + anti-IFNAR treated animals. However, the Tamiflu + anti-CD8 treated animals showed only a mild decrease in surfactant genes at 6 dpi compared to uninfected controls with no increase in the loss at 8 dpi (Fig. 7A). This was accompanied by reduced viral clearance with a high level of viral RNA still present on 12 dpi (Figs. 7B and S9A-S9E). These data suggest that Tamiflu can diminish viral damage while not entirely preventing continued epithelial cell infection and that CD8^+^ effector T cells contribute to loss of lung capacity at late times by killing these infected cells. Multiplex immunofluorescence imaging of lungs from animals treated with the combination of anti-CD8 antibody and Tamiflu revealed more staining for the virus at 8 dpi (p-value = 0.027) (Fig. S9F) associated with an increased number of pSPC-producing cells that were positive for IAV (p-value = 0.001) (Figs. 7C-7E). There was no significant difference in the absolute number of AT2 cells in the tissue section between mice treated with Tamiflu alone and the combination with anti-CD8 antibody (p = 0.12). However, we found a distinct distribution of pSPC^+^ cells; animals treated with Tamiflu alone showed regional loss of AT2 cells while Tamiflu together with anti-CD8 antibody treatment significantly preserved AT2 cell density and distribution throughout the tissue (p-value = 0.015) (Figs. 7F and G). Altogether, our results indicate that in combination with suboptimal anti-viral drug therapy, inhibition of type I IFN signaling or depletion of CD8^+^ T effector cells improves survival via distinct mechanisms, but with both acting in sum to rebalance repair vs. immune damage to favor lung function (Fig. 7H).

**Fig. 7.**
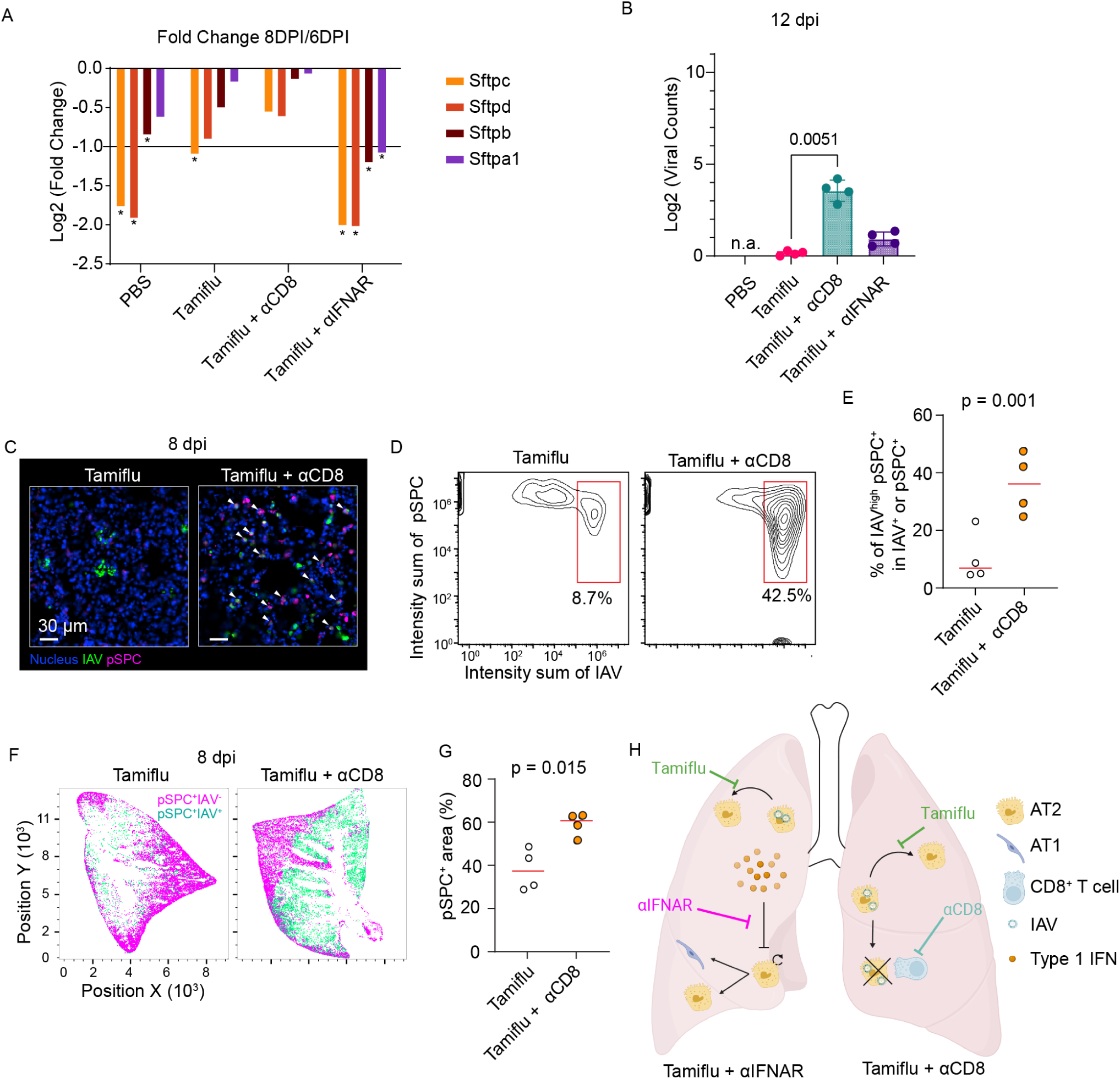
Combination strategies that target CD8^+^ T cells in addition to viral-mediated damage prevent AT2 cell loss. (A) Log2 fold change of 4 surfactant genes in each condition between 6 and 8 dpi. Asterix represents if the FDR from the edgeR glmLRT test was < 0.05 in the comparison. (B) Viral RNA counts at 12 dpi. n.a. is ‘not applicable’ since there were no survivors to 12 dpi in the PBS condition. p-values calculated from a Kruskal-Wallis test. (C) Immunofluorescence images of AT2 cells in lungs from PR8 infected mice treated with Tamiflu or Tamiflu + anti-CD8 antibody starting on 4 dpi. The lungs were harvested on 8 dpi. Low magnified views are shown in Fig. S8F. White arrows represent IAV^+^pSPC^+^ cells. (D and E) Contour plots of AT2 population by histocytometry (D) and the quantification (E). (F and G) Histocytometric visualization of pSPC^+^ IAV^−^ and pSPC^+^IAV^+^ populations (F) and the quantification of covered area of tissue with pSPC^+^ cells (G). Each dot represents an individual animal, and the red bars represent the median. Data are representative of two independent experiments except for (A and B); data from one experiment. n = 3-4 mice/group. (H) The diagram of the distinct mechanisms following treatment that rebalance pulmonary repair versus damage.

## DISCUSSION

Finding treatment strategies that prevent death when administered late in the course of severe viral pneumonia has proven difficult. Anti-viral drugs are the first and only choice for hospitalized patients beyond supportive care and early treatment with such agents is highly recommended (*15*). Here we have applied multiple analytical technologies to uncover the factors preventing effective treatment with either anti-inflammatory or anti-viral agents when started late in disease. Monotherapy that depletes neutrophils or prevents direct viral cytopathic effect enhanced survival in a lethal influenza infection model, but only if given very early. These findings indicated that the damage arising shortly after disease onset overwhelms the repair capacity of the tissue and passes a tipping point of minimal pulmonary function, with homeostasis difficult to regain, consistent with the poor prognosis of patients on ventilation. The corollary of such an interpretation is that interventions that enhance repair sufficiently to overcome this damage-induced deficit, when used with treatments that attenuate further direct viral tissue destruction, might be successful and indeed, that is what we found. Combining late, suboptimal anti-viral therapy with either blockade of type I IFN signaling to promote epithelial repair or limiting additional immune damage from late arriving effector CD8^+^ T cells to permit repair to overcome the existing physiological deficit saved animals from otherwise lethal infection. These findings show that later in viral pneumonias and possibly also in the case of various forms of ARDS, recovery will be favored if there is a rebalancing of immune- and viral-mediated damage vs. tissue repair.

Two lines of evidence suggested to us that type I interferons might be key to rebalancing damage and repair. First, while type I IFNs are a crucial component of host immunity with anti-viral and immune regulatory functions, these cytokines have been recognized to be pathogenic in pulmonary infectious diseases such as influenza, SARS-CoV, and SARS-CoV2 infections (*13, 33, 34*). Previous studies have highlighted that the timing of type I IFN production is critical to the protective vs. pathogenic influence of this innate response in SARS-CoV infection and MERS-CoV infection models (*18, 35*). Early administration (≤5 days after hospitalization) of IFN-α2b was associated with reduced mortality of hospitalized patients with COVID-19, whereas late administration was associated with increased mortality (*36*). Our transcriptome approach confirmed strong and sustained upregulation of interferon-stimulated genes (ISGs) (Fig. S5A) in this lethal influenza infection model. These published data, our RNAseq results, and the findings of Wack and colleagues showing that type I and type Ⅲ IFN inhibit AT2 cell proliferation and action as a source of stem cells for repair (*19*) all served as an impetus for testing late treatment of anti-IFNAR with Tamiflu as a therapy.

Beyond the salutary effects on survival, mechanistic studies using imaging and sequencing revealed that anti-IFNAR plus Tamiflu had a positive effect on the AT2 population. Other cell types such as KRT5^+^ KRT14^+^ p63^+^ basal cells(*37, 38*) also serve as sources of regeneration following lung damage. Our transcriptome analysis showed upregulation of *Krt5*, *Krt14*, and *Trp63* genes in the animals treated with Tamiflu + anti-IFNAR at 8 dpi (Fig. 6D), suggesting the induction of KRT5^+^ KRT14^+^ p63^+^ basal cells also contributed to improved survival. In addition, mesenchymal cells were also uniquely increased under these conditions (Fig. S7B). This cell type provides fibroblast growth factor (FGF) ligands that act on epithelial stem cells to promote airway regeneration and on AT2 cells to promote alveolar repair (*39*). In line with this, our histological data showed a more robust epithelial layer present at 8 dpi (Figs. S8C and S8D), along with activation of STAT3 signaling that can activate mesenchymal cells via the BNDF-TrkB pathway (*31*). The effect on mesenchymal cells became significant in the Tamiflu alone condition by 12 dpi (Fig. S8B), indicating that inhibition of type I IFN signaling accelerated the repair by activating mesenchymal and basal cells at an earlier stage of infection. This emphasis on the need for effective function of AT2 pneumocytes and associated stromal elements supporting epithelial repair in surviving severe viral infection is consistent with the observations made using scRNAseq of samples from patients dying of COVID-19 that revealed stress of the AT2 population as a major signal (*26*).

The second successful approach to preventing lethality when starting treatment well into the course of influenza infection involved depletion of CD8^+^ T cells. This strategy emerged from several considerations: first, the known capacity of cytotoxic CD8^+^ lymphocytes to clear virus by killing infected cells and preventing the further spread of the virus (*40, 41*), second, the obvious loss of cells that may have some remaining function even if infected, as well as the damage to bystander cells in the vicinity of the infected target cells by both cell-damaging mediators released from the cytotoxic effectors along with local stimulation of myeloid cells to produce cytotoxic cytokines, and third, the reported association of a delay in CD8^+^ T cell responses with worse outcome in COVID-19 infection (*42*). Together with a substantial literature on immunopathology mediated by CD8^+^ T cells in infected animals, we postulated that the damage caused by the delayed activity of adaptive immunity might exceed the benefit to the host of contributing to virus clearance and thus, depletion of these effectors might promote enhanced survival. This is what was observed, at the price of permitting continued viral replication that was attenuated by the simultaneous use of Tamiflu, even though this anti-viral drug was incapable on its own of salvaging the same number of animals as the combined treatment. Our sequencing studies showed that a high viral load distinguished animals treated with anti-CD8 from those given anti-IFNAR, making clear that two different mechanisms accounted for the benefit of these late treatment regimes. In the latter case, repair per se is not enhanced based on the RNAseq data (Figs. 6A, 6B, and S8A), but suppression of killing of infected cells also rebalances the rate of repair vs. additional cell death to favor survival.

Although none of the systematically tested 50 regimens in our *in vivo* screening approach were successful other than a marginal positive effect of agents targeting neutrophils, this does not necessarily exclude the potential efficacy of those treatments, especially in combination with effective anti-viral therapy. Indeed, our results suggest that the current recommendation against use of Paxlovid more than 5 days after symptoms begin for COVID-19 or 3 days in the case of influenza and Tamiflu may not maximize use of these drugs. These agents, as seen with Tamiflu late in influenza infection in our model system, could synergize with additional treatments to ameliorate severe disease.

In summary, here we provide evidence for a ‘tipping point’ model of lethal viral pulmonary infection, with early innate inflammatory events poising the animals for death even if further inflammatory damage is inhibited. Using anti-viral drugs to limit continued viral spread, we show that by improving the capacity to repair this existing damage or attenuating late adaptive immune damage to the pulmonary epithelium to allow ongoing repair to reach an effective level, we can overcome this early programming for lethality. Our approach highlights the value of focusing on the temporal evolution of distinct causes of tissue damage in viral ARDS and of the effects of the evolving immune response on repair processes to design more effective therapies when early use of potent anti-virals has not occurred. These findings provide a rationale for the use of anti-viral drugs late in the course of severe disease even though current recommendations are for their use only shortly after symptoms begin, markedly extending the potential utility of these agents when used in combination with other treatments and providing a basis for future clinical studies aimed at ameliorating severe influenza when anti-viral treatments are no longer effective on their own.

## MATERIALS AND METHODS

### Animals, Infections, and tissue collection

8 to 12-week-old male C57BL/6 mice were infected with different doses of H1N1 influenza A viruses via the oropharyngeal route under isoflurane anesthesia. Infectious doses were supplied in a volume of 50 μL sterile 0.9 % sodium chloride (Farris Laboratories). Drugs or antibodies were injected intraperitoneally in 200 μL volumes. Peripheral blood, the lung-draining lymph node, bone marrow, and lungs were harvested at various time points post-infection. For histology, left lobes were fixed with BD CytoFix/CytoPerm (BD Biosciences) diluted in phosphate-buffered saline (PBS) solution (1:4) for 1 day at 4 °C. Following fixation, tissues were washed briefly three times (5 min per wash) in PBS and incubated in 30% sucrose for 1 day at 4 °C before embedding in OCT compound (Sakura Finetek, Cat. #4583).

All mice were maintained in specific pathogen-free conditions at an Association for Assessment and Accreditation of Laboratory Animal Care accredited animal facility at the National Institute of Allergy and Infectious Diseases (NIAID). All procedures were approved by the NIAID Animal Care and Use Committee (NIH) under ASP LISB-4E.

### Virus

All studies used influenza H1N1 A/PR/8/34 (*43, 44*) seeds (a kind gift from J.R. Bennink, NIAID, NIH) propagated in EX-CELL MDCK Serum-Free Medium for MDCK Cells (Millipore Sigma, Cat. #14581C-1000ML).

### Drug treatment

All drugs and the regimens used in this study are listed in tables S1 and S2.

### Immunostaining

Frozen lung tissues were cut at 30 μm on a CM3050S cryostat (Leica) and adhered to Superfrost Plus slides (VWR, Cat. # 48311-703) coated with 15 μL of chrome alum-gelatin adhesive (Newcomer Supply, Cat. #1033A). Frozen sections were washed with PBS and then made permeable and blocked in PBS containing 0.3% Triton X-100 (Sigma, Cat. # T8787-100ML) and 1% BSA (Sigma-Aldrich, Cat. # 10735086001), and 1% mouse Fc block (BD Bioscience, Cat. #553142) followed by staining with antibodies diluted in blocking buffer. The antibodies used in this study are listed in table S3. After staining, slides were mounted with Fluormount G (Southern Biotech, Cat. #0100-01) and examined on a Leica Thunder Imager or TCS SP8 confocal microscope.

### Iterative bleaching and extend multiplexity (IBEX)

Iterative staining was performed in the principle of the IBEX protocol (*45, 46*). In brief, followed by the procedure of immunostaining above, fluorophore inactivation was performed with 1 mg/mL of LiBH_4_ for 15 minutes to inactivate all fluorophores except for Hoechst after image acquisition. Samples were further stained with the antibodies for the second cycle of imaging. Image acquisition was carried out on the same microscope with the same X and Y positions. Z plane was adjusted to match the previous round of image with the focus map function in LAS X Software (Leica Microsystems). Image registration was performed with SimpleITK as previously described (*45, 46*).

### Image acquisition

Images were acquired using an inverted Leica SP8 confocal microscope equipped with a 20 ⨉ objective (NA 0.95), 4 HyD and 1 PMT detectors, a white light laser that produces a continuous spectral output between 470 and 670 nm as well as 405, 685, and 730 nm lasers or Leica THUNDER Imager Tissue equipped with a 20⨉ objective (NA 0.75) (Leica, Cat #11506343), DFC9000 GTC cMOS camera (Leica), LED 8 Fluorescent light source (Leica), CYR71010 filter cube (Leica, Cat. #11525416), and DFT51010 filter cube (Leica, Cat. #11525418). Panels consisted of antibodies conjugated to the following fluorophores and dyes: Hoechst, eFluor (eF) 450, Alexa Fluor (AF) 488, AF532, PE, AF594, AF647, AF700, AF750, DyeLight 755. Images were converted to .ims file with Imaris File Converter (Oxford Instruments) and visualized by Imaris software (Oxford Instruments). Gaussian filters were applied to denoise.

### Image processing and analysis

Images were converted to .ims file with Imaris File Converter (Oxford Instruments) and gaussian filters were applied to all images with Imaris. For cell segmentation and quantification, .ims files were converted to OME-tiff format. The Tissuenet model in Cellpose 2.0 (*47, 48*), further trained with our dataset, was used to segment cells. Mean intensity of each marker and X, Y coordinates were extracted by 3D ImageJ Suite for FIJI (*49*). Extracted values were combined into a single csv file and analyzed by histocytometry as previously described (*50*). To quantify IAV-NP^high^ pSPC^+^ cells, a composite image of IAV-NP and pSPC was created by adding those channels with Imaris channel arithmetic function after subtracting background. Intensity sum of IAV-NP and pSPC were exported and analyzed by Flowjo software as described previously (*50*). For the coverage of pSPC^+^ cells in the tissue, total nucleus was identified with the Imaris Spots function with the estimated diameter was 6 μm and their X and Y coordinates were exported as a csv file. pSPC^+^ cells were gated by Flowjo based on the intensity sum, and their X and Y coordinates were exported as a csv file. A grid of 100 × 100 μm was artificially created on the image. If the number of total nuclei exceeds 20, the number of pSPC^+^ cells is counted on the region to exclude empty regions out of the tissue. If the number of pSPC^+^ cells is greater that 5% of the total nucleus in the regions, those regions are considered as covered regions. The code was written in Python 3.9.13.

### RNA preparation for RNA-Seq

Right inferior lobe was immediately transferred into 1 mL Trizol (Thermo, Cat. #15596018), snap-frozen in liquid nitrogen, and kept at −80 °C. RNA extraction was performed by QIAGEN Nucleic Acid Isolation Service (QIAGEN, Germany). RNA integrity number (RIN) was determined with Agilent RNA 6000 Nano Kit (Agilent, Cat. # 5067-1511) for quality check and all samples were above 8.5.

### Next Generation Sequencing

Sequencing libraries were made from 1 µg total RNA using the TruSeq Stranded mRNA kit, according to the manufacturer’s guide (Illumina, San Diego, CA). To obtain reads per sample, two microliters of each purified library was combined into a single pool, quantified using the Kapa Library Quantification kit (Roche Sequencing Solutions, Pleasanton, CA), diluted to 10 pM, and sequenced as 2 × 151 bp reads on the MiSeq instrument using the Nano kit, V2 (Illumina, San Diega, CA). A second library pool was created where the volume of each library was calculated based on the read distribution from the MiSeq run to create a normalized pool for deep sequencing. The normalized pool was quantified using the Kapa Library Quantification kit, diluted to 1.8 pM, and sequenced as 2 × 151 bp paired-end reads on three NextSeq 550 instrument runs using the High Output v2.5, 300 cycle sequencing kit. A total depth of 1.04 billion paired-end reads with an average of 15.3 million paired-end reads per sample was generated.

### RNA-Seq Data Processing

Following sequencing, samples were aligned to the mouse genome (GRcm39) appended to contain the influenza virus genome (GCF_000865725.1) to account for viral RNA reads. Alignment to the genome was done using STAR v2.7.10 (*51*) and counts were generated using featureCounts in the SubRead package v2.0.4 (*52*). The count tables were read into R v4.2.1. Data normalization and differential expression analysis were performed using edgeR v3.40.2 (*53*). Pathway analysis was performed using the fgsea package v1.24.0 (*54*) and the c2cp gene set from the MSigDB (*55*). Pathways were clustered into groups using a Euclidian distance metric on the Jaccard index of similarity calculation between genes included in the pathway list. Cell type deconvolution from RNA-Seq data was performed using the MuSIC package v1.0.0 (*56*) in R and using the naïve lung single cell data form (*57*) as a background. Results were exported to GraphPad/Prism v10.1.1 for display only.

### Quantification and statistical analysis

Statistical tests performed for a given analysis are denoted in the figure legends. For this manuscript, a p-value less than 0.05 was considered as a significance threshold. All statistical tests were performed in R or GraphPad/Prism v10.

## ACKNOWLEDGMENTS

We thank J.R. Bennink for providing PR8 influenza A virus. This research was supported by the Division of Intramural Research of the National Institute of Allergy and Infectious Diseases, NIH. RNA-Seq was performed by Genomic Research Section, Research technologies Branch, NIAID. We thank Calithera Biosciences for providing CB-280. We thank U. Undersson for providing anti-HMGB1 antibody.

## AUTHOR CONTRIBUTIONS

Conceptualization, R.N.G.; Methodology, H.I., F.L.R., T.Z.V., C.J.C., A.G., E.S.; Validation, H.I., F.L.R., C.J.C., A.G.; Formal Analysis, H.I., F.L.R., E.S.; Investigation, H.I., F.L.R.; Data Curation, H.I., E.S.; Writing – Original Draft, H.I., E.S.; Writing – Review & Editing, R.N.G.; Supervision, R.N.G.; Project Administration, H.I., R.N.G.; Funding Acquisition, R.N.G.

## DATA AVAILABLITITY STATEMENT

Any information required to reanalyze the data reported in this paper is available from the lead contact upon request.

## Supplementary Materials

### Materials and Methods

#### Serum cytokine measurement

Blood obtained by cardiac puncture was placed in Microvette 500 Serum Gel tubes (Sarstedt, 20.1344) and allowed to clot before analysis with the Lunaris Mouse 12-Plex Cytokine Kit (Ayoxxa Biosystems).

#### qRT-PCR

Lung samples were collected into tubes containing Trizol (Invitrogen) and manually disassociated in the Trizol followed by processing on direct-zol columns (Zymo Research). RNA concentration was determined on a nanodrop (Thermo Fischer Scientific). cDNA was generated using the SuperScript IV VILO master mix (Invitrogen) and 100 ng/µL of RNA following a program of 25 degrees for 10 minutes, 50 degrees for 10 minutes, and 85 degrees for 5 minutes. cDNA was added to a master mix containing Platinum Quantitative PCR Supermix UDG with ROX (Invitrogen) and the TaqMan (Invitrogen) primer/probe set for influenza A or *Hprt*. Readout was performed in triplicate for each sample using a 15 μL reaction volume in a 384-well plate on the Quant Studio 6 (Thermo Fischer). The viral ct values were normalized against *Hprt* using the ΔΔ method and the change compared to the average of the controls was calculated.

#### Flow cytometry

For the lungs, the right cranial lobes were used for all experiments. Harvested lungs were chopped in 2 mL of a mixture of liberase TM (100 µg/mL) (Sigma, Cat. #5401127001) and DNaseI (100 μg/mL) (Roche, Ct. #04716728001) in RPMI-1640 (Thermo Fisher Scientific, Cat. #11875093) and transferred to gentleMACS C Tubes (Miltenyi Biotech, Cat. #130-093-237). Samples were homogenized by 2 cycles of mechanical dissociation with a gentleMACS Dissociator (Miltenyi Biotech, Cat. #130-093-235) and incubated at 37 °C for 30 minutes for enzymatic digestion in between. Single-cell suspensions were obtained by adding EDTA at a final concentration of 20 mM to stop the enzymatic reaction and passing through a 70 μm nylon mesh. Cells were washed in PBS with 2% FBS, 5 mM EDTA (hereafter FACS buffer) twice. The lung-draining lymph nodes were harvested and digested in 500 μL of RPMI with liberase TM (200 μg/mL) at 37 °C for 30 minutes. Single-cell suspensions were obtained by adding EDTA at a final concentration of 20 mM to stop the enzymatic reaction and passing through a 70 μm nylon mesh. Cells were washed in FACS buffer twice. Bone marrow cells were flushed from the femora and tibiae. Suspensions were passed through a 70 μm nylon mesh. Peripheral blood was collected from the inferior vena cava. BD Pharm Lyse Lysing Buffer (BD Biosciences, Cat. #555899) was used to lyse red blood cells. Cells in FACS buffer were blocked with anti-CD16/CD32 (BioLegend, Cat. #101302) and then were incubated with fluorochrome-conjugated antibodies with LIVE/DEAD fixable dead cell stain dye (Thermo Fisher Scientific, Cat. # 423108). Staining and washing were performed at 4 °C. CountBright Absolute Counting Beads (Thermo Fisher Scientific, Cat. # C36950) were added into cell suspension before analysis. Cells were analyzed on an LSR Fortessa flow cytometer (BD Biosciences) and data were analyzed with FlowJo software (BD).

#### Microscope configuration

Images were acquired using an inverted Leica TCS SP8 X confocal microscope equipped with a 20⨉ objective (NA 0.75), 4 HyD and 1 PMT detectors, a white light laser that produces a continuous spectral output between 470 and 670 nm as well as 405, 685, and 730 nm lasers or Leica THUNDER Imager Tissue equipped with a 20⨉ objective (NA 0.75) (Leica, Cat #11506343), DFC9000 GTC cMOS camera (Leica), LED 8 Fluorescent light source (Leica), CYR71010 filter cube (Leica). Images were captured at 12-bit, with a line average of 3, and 1024×1024-pixel format for the confocal imaging or 8-bit depth or 16-bit, 2048×2048-pixel format for the Thunder imaging. Tiling images were acquired and merged using the LAS X software.

#### Tissue clearing

Volumetric imaging with optically cleared samples were performed as described previously with slight modification. Briefly, frozen lung samples were sectioned at 500 μm on a CM3050S cryostat. The samples were hydrated and washed with PBS to remove OCT in a 24-well plate. Samples were incubated for at least 12 hours in BD Perm/Wash Buffer (BD Bioscience) containing 1% mouse Fc block (BD Bioscience, Cat. #553142) and stained with titrated antibodies in BD Perm/Wash Buffer (BD Bioscience, Cat. #554723) containing 1% Fc block for 24 hours at room temperature on a shaker. Stained samples were washed with BD Perm/Wash Buffer three times for at least 20 minutes at room temperature on a shaker. Samples were transferred on a slide with two silicon isolators (Grace BioLabs, Cat. #664407) and treated with 200 μL of Ce3D medium [1.82 g Histodenz (Millipore Sigma, Cat. #D2158-100G, 0.1% triton, and 0.5% thioglycerol (Millipore Sigma, Cat. #M1753) per 1 mL 40% N-methylacetamide (Millipore Sigma, Cat. #M26305–500G) in PBS] inside a chemical fume hood and sealed with a cover slip (Electron Microscopy Sciences, Cat. #63766-01) and incubated at room temperature on a shaker overnight. After removing the old Ce3D, cleared samples were mounted with 40 μL of new Ce3D and sealed with a coverslip with two SecureSeal Imaging Spacers (Grace Bio-Labs, Cat. #654002).

**Fig. S1.**
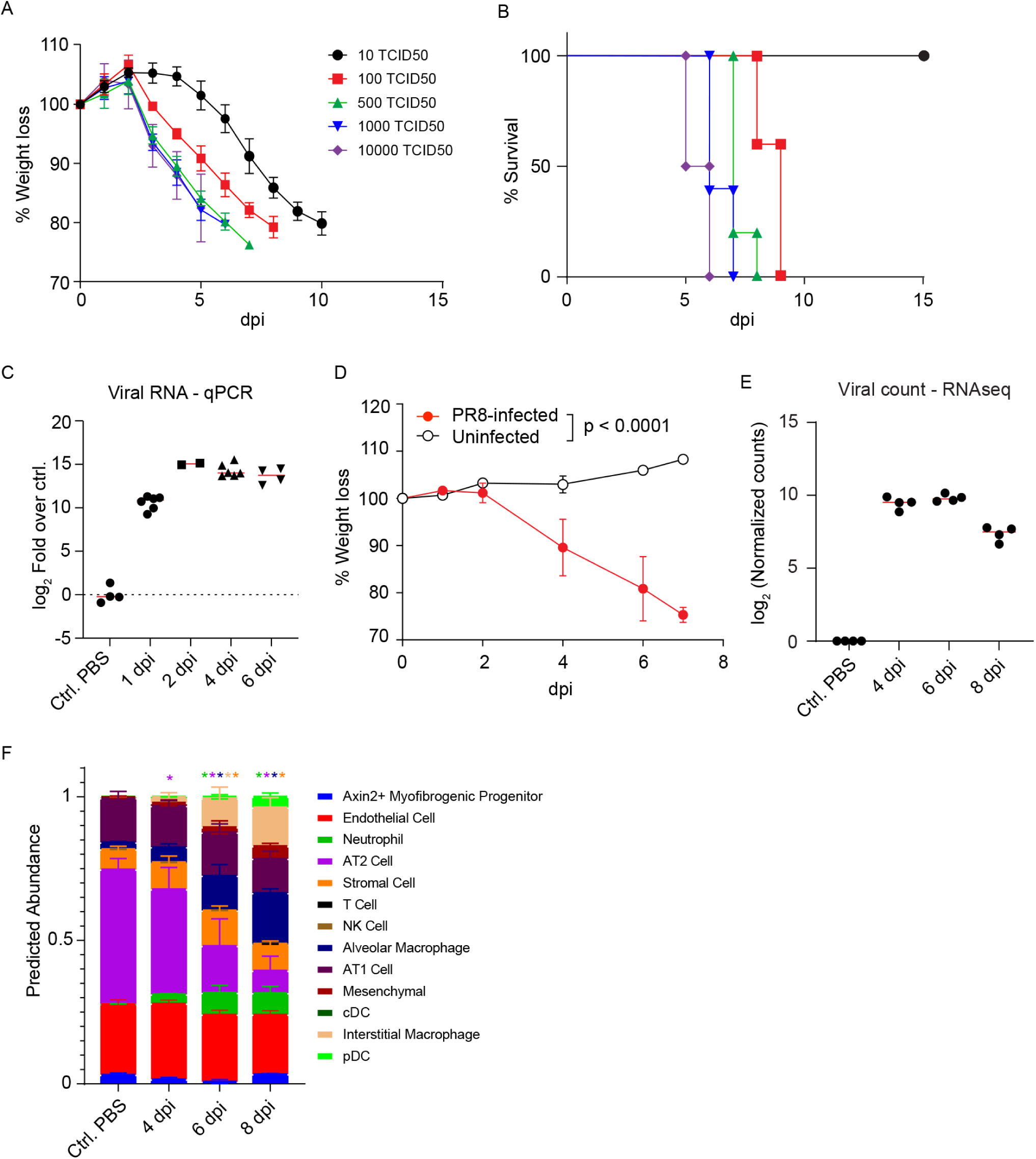
(A and B) Weight and survival curves for various doses of PR8. Data are from one cohort with n = 4 or 5. (C and D) The time course of viral titer and body weight at 100 TCID50. Each dot represents an individual animal in (C). Bars represent mean ± standard error of the mean (SEM). Data are from one cohort with n=7 (PR8-infected) and n=5 (Uninfected) mice/group in (D). P-values were determined by a two-way ANOVA. (E) Log 2 normalized counts of viral RNA reads from RNA-seq. Each point represents a single animal and the bar represents the geometric mean of the timepoint. (F) Cell-type deconvolution results from RNA-Seq. Each colored bar is the mean representation of a given cell type with the standard deviation from all the samples. The colored asterisk at the top denotes if a given cell type was significantly different than the control sample. A 2-way ANOVA was used to determine significance. The y-axis is the predicted percent abundance of a given cell type from the deconvolution.

**Fig. S2.**
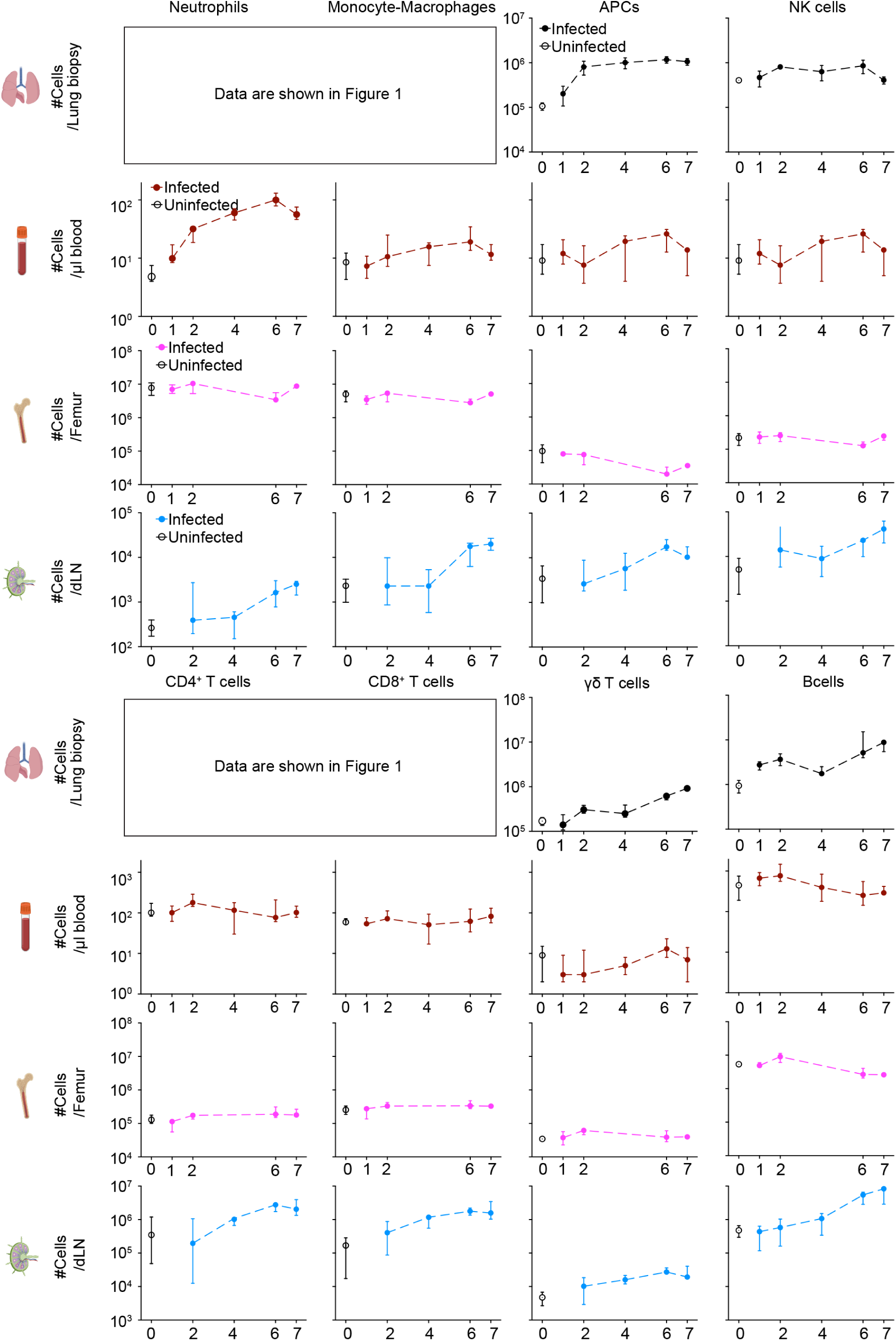
Flow cytometric analysis of absolute cell numbers in the lungs, peripheral blood, bone marrow, and LDLN samples, related to Fig. 1E and 1F. Data are representative of at least two independent experiments with n = 7 mice/group. Bars represent mean ± SEM.

**Fig. S3.**
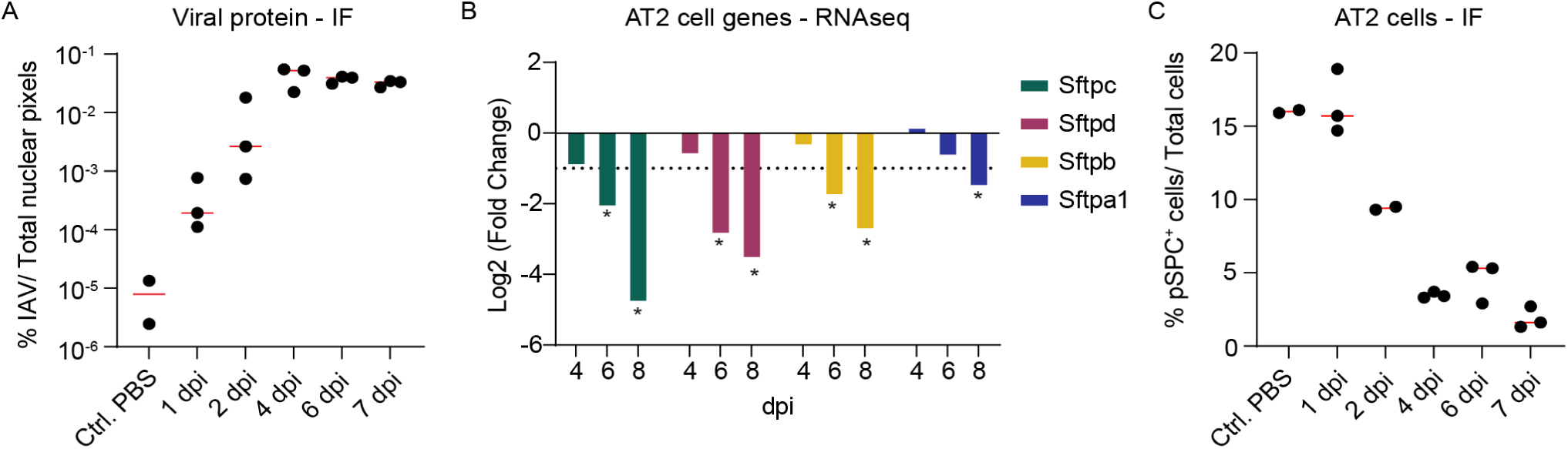
(A) The time course of viral protein expression quantified at the pixel level by immunofluorescence. (B) Log 2-fold change of the surfactant genes from the RNA-Seq data for each day compared to uninfected controls. The dashed line represents a fold change greater than 2. An asterisk indicates that the fold change was significant as determined by the glmLRT test in edgeR DE results. (C) The time course of pSPC^+^ cell number determined by immunofluorescence imaging. Each dot represents an individual animal and red bars represent the median. IF means immunofluorescence. Each dot represents an individual animal and data are from one cohort.

**Fig. S4.**
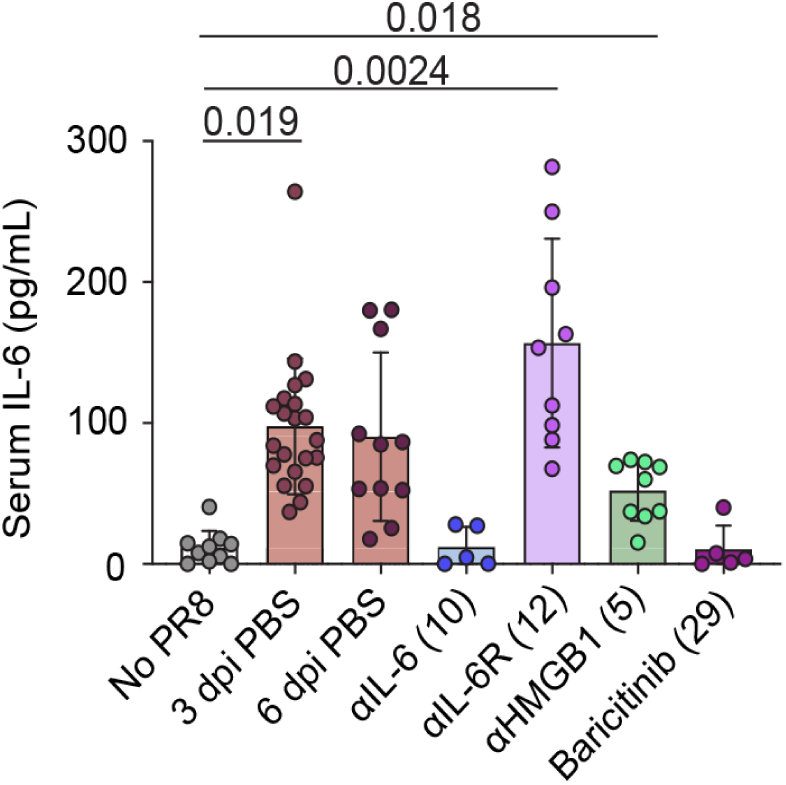
Representative analysis of serum cytokines to assess drug efficacy. Sera were collected at 6 dpi. Each dot represents an individual animal and bars represent mean ± SEM. The numbers in parentheses correspond with the treatment regimens as shown in table S2. Data are pooled from 1 – 4 independent experiments. Adjusted p-values are determined from a Kruskal-Wallis test and shown on top.

**Fig. S5.**
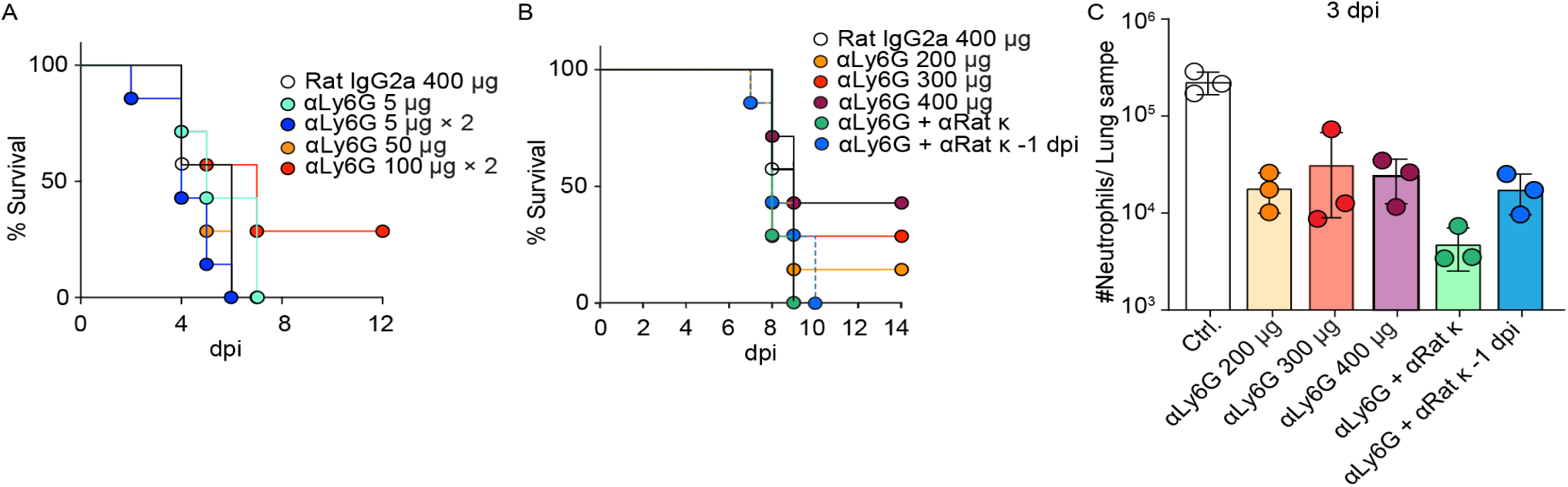
(A and B) Survival after PR8 infection upon treatment as indicated. Data are from one cohort with n=7 or 8 mice/group. (C) Flow cytometric analysis of neutrophils in the lung biopsy samples. Samples were harvested on 3 dpi. Data are from one cohort with n=3 mice/group.

**Fig. S6.**
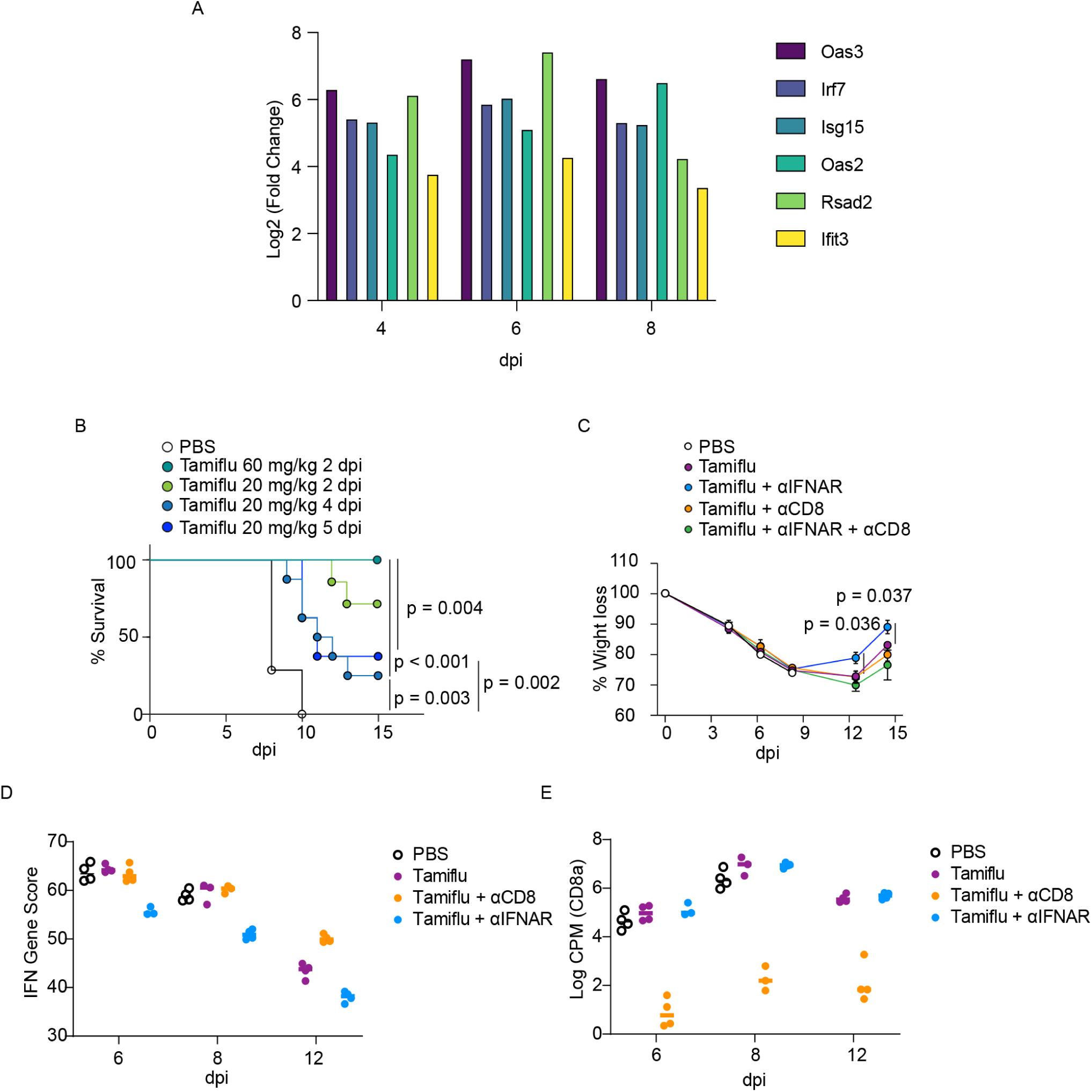
(A) Log 2-fold change of 5 different interferon-stimulated genes over the course of infection in untreated animals as compared to uninfected controls. All differences between untreated and uninfected animals were significant based on the results of the glmLRT differential expression test in edgeR. (B) Survival after PR8 infection and the indicated Tamiflu treatment. Data are from one cohort with n=7-8 mice/group. (C) Percent weight loss after infection with indicated treatments. Data are representative of two independent experiments. n = 8 mice/group. Bars represent mean ± SEM. (D) Gene module score of the ISG expression for 6 type I interferon responsive genes in the different conditions across the time course. Each point represents a single animal. (E) Log counts per million of normalized expression data of the CD8a gene from the RNA-Seq data. Each point represents a single animal. Data are from one cohort with n=4 mice/group (A, D, and E).

**Fig. S7.**
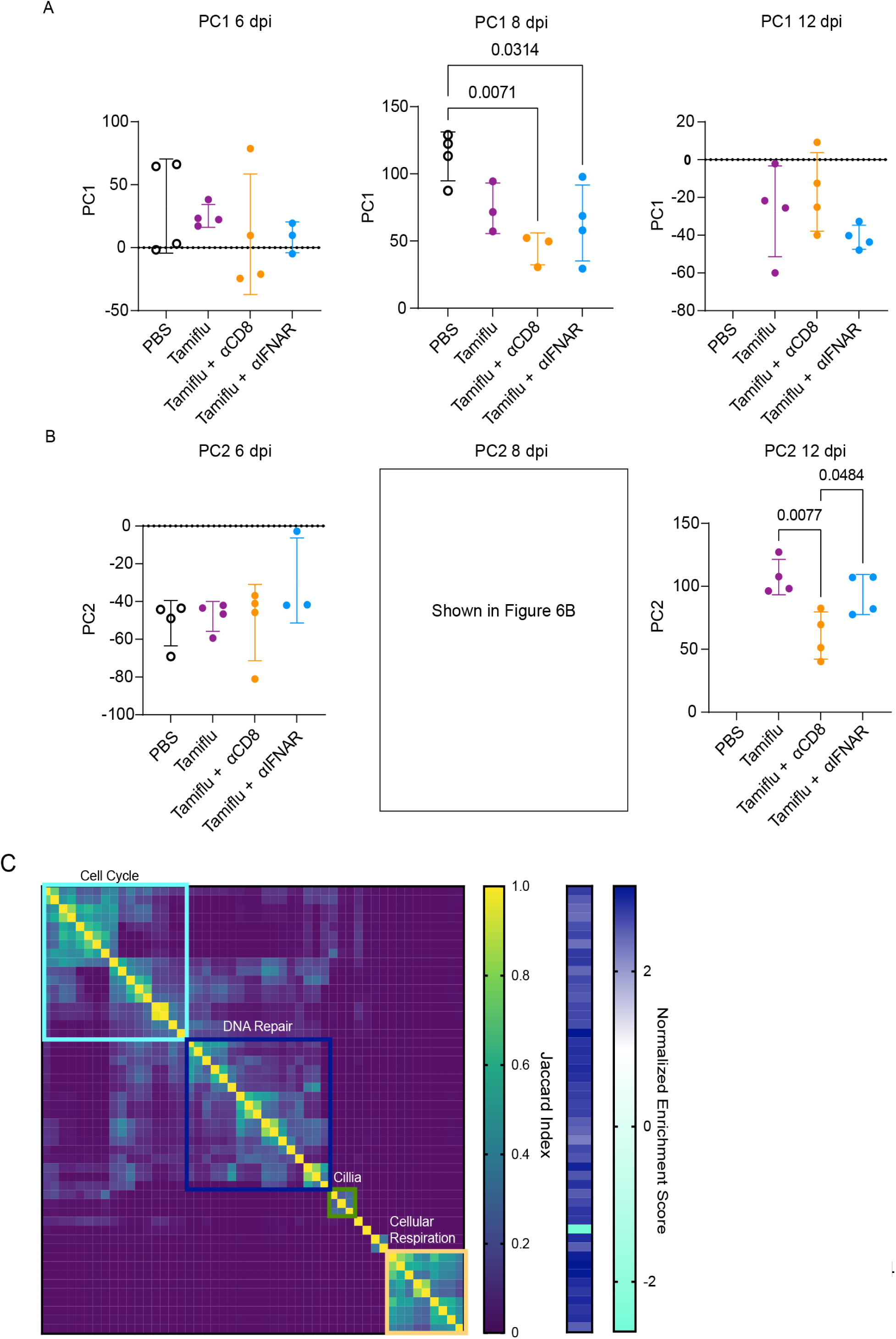
(A) Location of all samples in the principal component space across PC1 separated by the different conditions and the dpi. Each point is a single animal, bars represent the mean of the group and error bars are the standard deviation. Significant comparisons are shown as p-values from a one-way ANOVA test. (B) Same as (A) but along principal component 2. (C) Jaccard similarity index of the top 50 pathways along principal component 2 as determined by adjusted p-value. Heatmap shows similarity index between pathways, which are rows and columns. Groupings are determined by clustering of pathways using a Euclidian distance metric. To the right of the heatmap, the normalized enrichment score of the pathways along PC2. Data are from one cohort with n=4 mice/group.

**Fig. S8.**
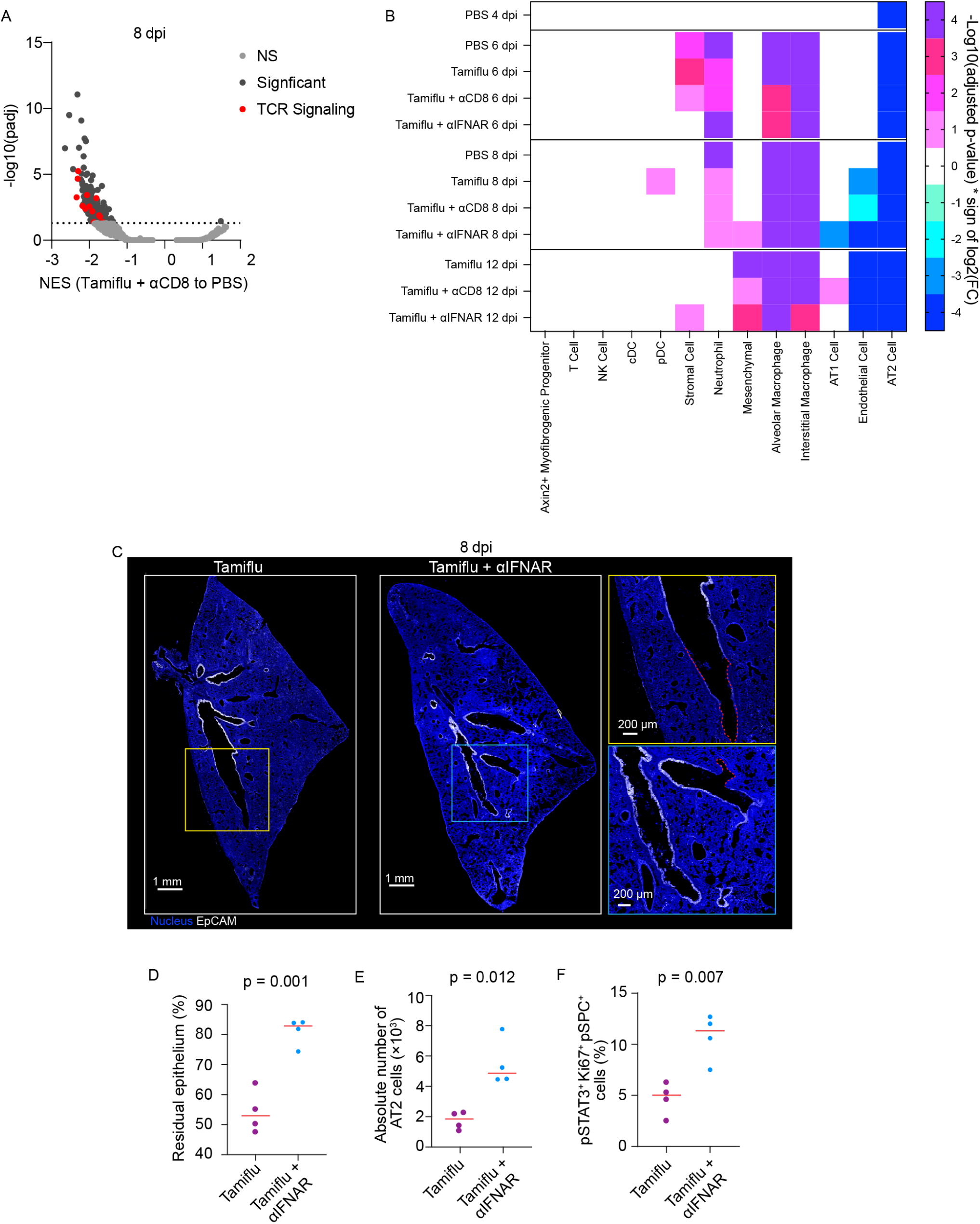
(A) Results of gene set enrichment analysis comparing data at 8 dpi for animals receiving Tamiflu + anti-CD8 or PBS. Pathways were grouped by Jaccard similarity and colored based on different groupings. NS is not significant. (B) Significance results from the cell type deconvolution of all samples compared to uninfected controls. Columns are cell types and rows are the different conditions. Colors are the −log10(p-value) multiplied by the sign of the log2 fold change. P-values were determined by a two-way ANOVA. (C and D) Immunofluorescence images and quantification of bronchial epithelial cells in lungs from PR8 infected mice treated with Tamiflu or Tamiflu + anti-IFNAR1 antibody starting on 4 dpi. The lungs were harvested on 8 dpi. Magnified views of highlighted regions are shown on the right. Dotted lines show the region that lost bronchial epithelium. (E and F) Quantification of the number of AT2 cells (E) and percentage of pSTAT3^+^Ki67^+^ AT2 cells (F) in the lung section, related to Fig. 5C-5F. Each dot represents an individual animal, and the red bars represent the median. Data are from one (A and B) or two (C and D) cohorts with n=4 mice/group.

**Fig. S9.**
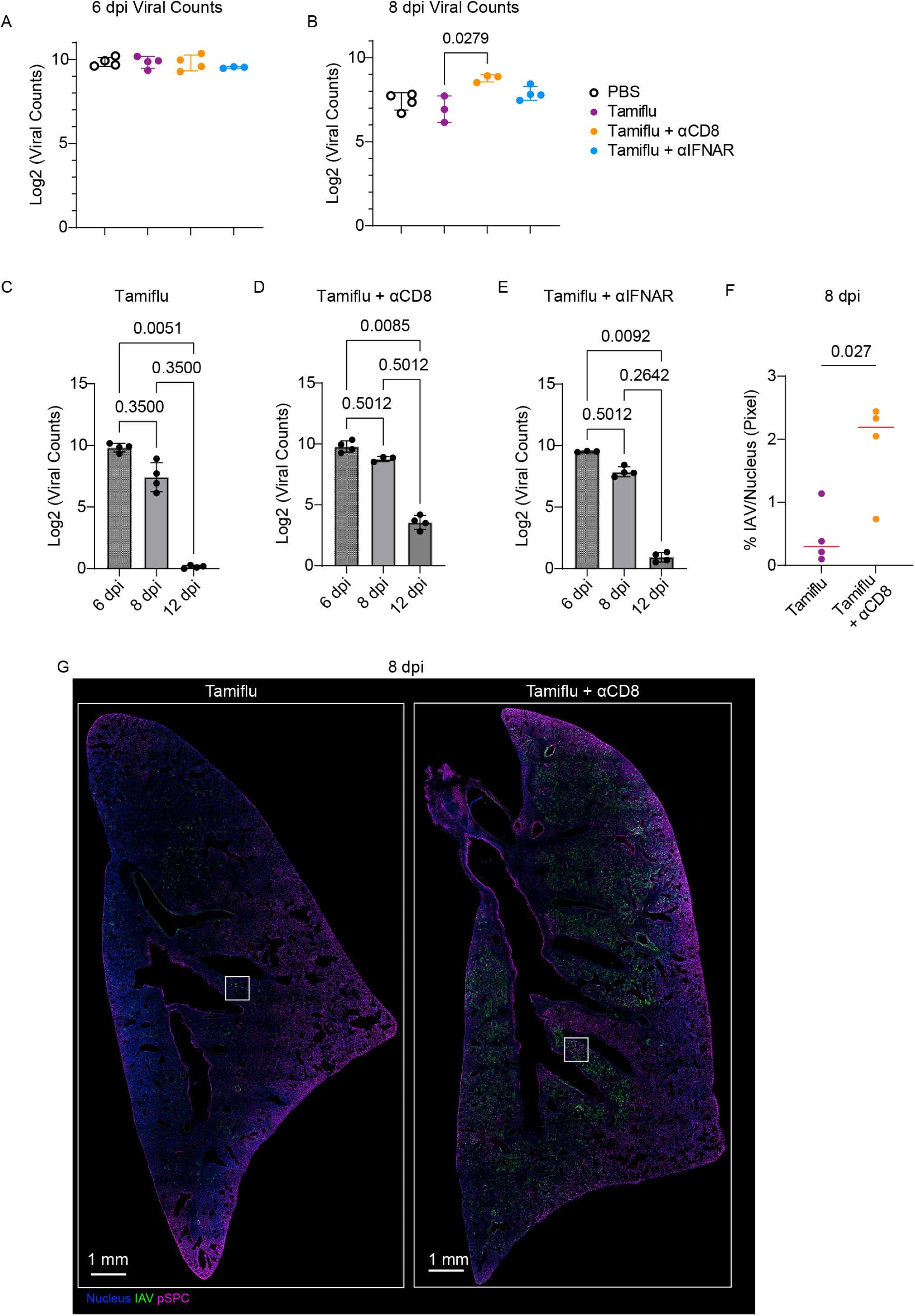
Log2 of normalized viral RNA counts from RNA-Seq data at 6 dpi and 8 dpi samples with indicated treatments (A and B), and from Tamiflu, Tamiflu + anti-CD8, and Tamiflu + anti-IFNAR treated samples with indicated dpi (C-E). P-values were calculated from a Kruskal-Wallis test. Data are from one cohort with n=4 mice/group. (F) Low magnified images of Fig. 7C. Immunofluorescence images of the lung infected PR8 and treated with Tamiflu + anti-CD8 antibody starting on 4 dpi. Tissues were harvested on 8 dpi. White squares indicate the regions shown in Fig. 7C.

**Table S1.** Drug list.

| Drug | Target | Dose | Route | Diluent | Vendor | Catalog Number |
| --- | --- | --- | --- | --- | --- | --- |
| Acalabrutinib | BTK | 25mg/kg | i.p. | 10%DMSO, 40% PEG300, 5% Tween-80, 45% saline (v/v) | Selleckchem | S8116 |
| Anti-CCL2 | CCL2 | 200 µg/mouse | i.p. | PBS | Bio X Cell | BE0185 |
| Anti-CD8 | CD8 | 250 µg/mouse | i.p. | PBS | Bio X Cell | BP0117 |
| Anti-CD40 | CD40 | 100 µg/mouse | i.p. | PBS | Bio X Cell | CP133 |
| Anti-HMGB1 | HMGB1 | 100 µg/mouse | i.p. | PBS | N/A | Gift from U. Undersson (58) |
| Anti-IFNAR1 | IFNAR1 | 500 g/mouse | i.p. | PBS | Bio X Cell | BE0241 |
| Anti-IL-6 | IL-6 | 200 µg/mouse | i.p. | PBS | Bio X Cell | BE0046 |
| Anti-IL6R | IL-6R | 200 µg/mouse | i.p. | PBS | Bio X Cell | BE0047 |
| Anti-Ly6G | Ly6G | 5-400 µg/mouse | i.p. | PBS | Bio X Cell | BE0075-1 |
| Anti-TNFα | TNFα | 200 µg/mouse | i.p. | PBS | Bio X Cell | BE0058 |
| Anti-PSLG-1 | PSLG-1 | 200 µg/mouse | i.p. | PBS | Bio X Cell | BE0186 |
| AZD-5069 | CXCR2 | 125 mg/kg | i.p. | 10%DMSO, 40% PEG300, 5% Tween-80, 45% saline (v/v) | MedChemExpress | HY-19855 |
| Baricitinib | JAK1/2 | 50 mg/kg | Sc. | 10%DMSO, 40% PEG300, 5% Tween-80, 45% saline (v/v) | MedChemExpress | HY-15315 |
| CB-280 | Arginase | 200 mg/kg | Sc. |  | Calithera | N/A |
| Colchicine | Inflammasome | 1 mg/kg | i.p. | Saline | Sigma | C9754 |
| Dexamethasone |  | 2.5 mg/kg | v | 10%DMSO, 40% PEG300, 5% Tween-80, 45% saline (v/v) | MedChemExpress | HY-14648 |
| Dipyridamole | Phosphodiesterase V | 1 mg/kg | i.p. | 10%DMSO, 40% PEG300, 5% Tween-80, 45% saline (v/v) | Sigma | PHR3772 |
| FGF-7 | FGFR2 | 0.5 µg/mouse | OPh. / i.p. | PBS | MedChemExpress | HY-P7230 |
| Ibrutinib | BTK | 25 mg/kg | i.p. | 10%DMSO, 40% PEG300, 5% Tween-80, 45% saline (v/v) | MedChemExpress | HY-10997 |
| Inosine Pranobex | Immunostimulant | 50 mg/kg | Sc. | PBS | Tokyo Chemical Industry | I1037 |
| Oseltamivir | influenza neuraminidase | 2-62.5 mg/kg | i.p. | PBS | MedChemExpress | HY-13318 |
| PAMAM dendrimer | Nuclear acid-containing debris | 20 mg/kg | i.p. | PBS | Sigma | 412422 |
| PMX205 | C5a receptor | 1 mg/kg | Sc. | Water | TOCRIS | 5196 |
| Ruxolitinib | JAK1/2 | 125 mg/kg | Oral | 10%DMSO, 40% PEG300,<br>5% Tween-80, 45% saline<br>(v/v) | Selleckchem | S1378 |
| Silvestat | Neutrophil elastase | 100 mg/kg | i.p. | 10%DMSO, 40% PEG300,<br>5% Tween-80, 45% saline<br>(v/v) | Selleckchem | S8136 |
| Topotecan<br>Hydrochloride | Topoisomerase I | 2 mg/kg | i.p. | 10%DMSO, 40% PEG300,<br>5% Tween-80, 45% saline<br>(v/v) | MedChemExpress | HY-13768A |
| Ziluton | 5-Lipoxygenase | 60 mg/kg | Sc. | 10%DMSO, 40% PEG300,<br>5% Tween-80, 45% saline<br>(v/v) | MedChemExpress | HY-14164 |
| Rat IgG2a<br>Isotype control. |  | 100 µg/mouse | i.p. | PBS | Bio X Cell | BE0089 |
| Anti-rat Kappa<br>immunoglobulin |  | 50 µg/mouse | i.p. | PBS | Bio X Cell | BE0122 |

**Table S2.** Treatment regimens.

|  | Treatments | Regimen | Dose | Route | N (Replicate) | Survived > 14 dpi |
| --- | --- | --- | --- | --- | --- | --- |
| 1 | Acalabrutinib | 2 – 7 dpi, once a day | 25 mg/kg | i.p. | 10 | 0 |
| 2 | Anti-CCL2 | 2, 4, and 6 dpi, once a day | 200 µg/mouse | i.p. | 10 | 0 |
| 3 | Anti-CD8 | 4 and 5 dpi | 250 µg/mouse | i.p. | 23 (3) | 0 |
| 4 | Anti-CD40 | 4 dpi | 100 µg/mouse | i.p. | 10 | 0 |
| 5 | Anti-HMGB1 | 4, 5, and 6 dpi, twice a day | 50 µg/mouse | i.p. | 10 | 0 |
| 6 | Anti-HMGB1 | 4, 5, and 6 dpi, once a day | 100 µg/mouse | i.p. | 10 | 0 |
| 7 | Anti-HMGB1 | 2 – 7 dpi, once a day | 50 µg/mouse | i.p. | 10 | 0 |
| 8 | Anti-IFNAR1 | 4, 6, and 8 dpi | 500 g/mouse | i.p. | 23 (3) | 0 |
| 9 | Anti-IL-6 | Dayly from 4 dpi, once a day | 200 µg/mouse | i.p. | 10 | 0 |
| 10 | Anti-IL-6 | 4, 5, and 6 dpi, once a day | 200 µg/mouse | i.p. | 10 | 0 |
| 11 | Anti-IL-6 | 2 – 7 dpi, once a day | 200 µg/mouse | i.p. | 10 | 0 |
| 12 | Anti-IL6R | 4, 5, and 6 dpi, once a day | 200 µg/mouse | i.p. | 10 | 0 |
| 13 | Anti-IL6R | 2 – 7 dpi, once a day | 200 µg/mouse | i.p. | 10 | 0 |
| 14 | Anti-Ly6G | 1 dpi, once a day | 5 µg/mouse | i.p. | 10 | 0 |
| 15 | Anti-Ly6G | 1 dpi, twice a day | 5 µg/mouse | i.p. | 10 | 0 |
| 16 | Anti-Ly6G | 1 dpi, once a day | 50 µg/mouse | i.p. | 10 | 0 |
| 17 | Anti-Ly6G | 1 dpi, twice a day | 100 µg/mouse | i.p. | 20 (2) | 2 |
| 18 | Anti-Ly6G | 1 dpi, once a day | 200 µg/mouse | i.p. | 10 | 1 |
| 19 | Anti-Ly6G | 1 dpi, once a day | 300 µg/mouse | i.p. | 10 | 2 |
| 20 | Anti-Ly6G | 1 dpi, once a day | 400 µg/mouse | i.p. | 27 (3) | 6 |
| 21 | Anti-Ly6G | 2 dpi, once a day | 400 µg/mouse | i.p. | 7 | 0 |
| 22 | Anti-Ly6G | 3 dpi, once a day | 400 µg/mouse | i.p. | 7 | 0 |
| 23 | Anti-Ly6G | 1 dpi, once a day | 25 µg/mouse | i.p. | 10 | 0 |
|  | Anti-rat Kappa immunoglobulin |  | 50 µg/mouse | i.p. |  |  |
| 24 | Anti-Ly6G | -1 dpi, once a day | 25 µg/mouse | i.p. | 10 | 0 |
|  | Anti-rat Kappa immunoglobulin |  | 50 µg/mouse | i.p. |  |  |
| 25 | Rat IgG2a Isotype control. | 1 dpi, twice a day | 100 µg/mouse | i.p. | 10 | 0 |
| 26 | Anti-TNFα | 2, 4, and 6 dpi, once a day | 200 µg/mouse | i.p. | 10 | 0 |
| 27 | Anti-PSLG-1 | 2 – 7 dpi, once a day | 200 µg/mouse | i.p. | 10 | 0 |
| 28 | AZD-5069 | 4, 5, 7 dpi, once a day | 100 mg/kg | i.p. | 10 | 0 |
| 29 | Baricitinib | 2 – 7 dpi, once a day | 50 mg/kg | Sc. | 10 | 0 |
| 30 | CB-280 | 2 – 7 dpi, once a day | 200 mg/kg | Sc. | 10 | 0 |
| 31 | Colchicine | 2 – 7 dpi, once a day | 1 mg/kg | i.p. | 10 | 0 |
| 32 | Colchicine | 2, 4, and 6 dpi, once a day | 0.5 mg/kg | i.p. | 10 | 0 |
| 33 | Dexamethasone | 2 – 7 dpi, once a day | 1 mg/kg | i.p. | 10 | 0 |
| 34 | Dipyridamole | 2 – 7 dpi, once a day | 2.5 mg/kg | v | 10 | 0 |
| 35 | FGF-7 | 4 dpi | 0.5 µg/mouse at 4 dpi | OPh. | 10 | 0 |
|  |  | 6 dpi | 5 µg/mouse at 6 dpi | i.p. |  |  |
| 36 | Ibrutinib | 2 – 7 dpi, once a day | 25 mg/kg | i.p. | 10 | 0 |
| 37 | Inosine Pranobex | 2 – 7 dpi, once a day | 50 mg/kg | Sc. | 10 | 0 |
| 38 | Oseltamivir | 2 - 6 dpi, twice a day for the first 3 days and once a day | 60 mg/kg | i.p. | 27 (3) | 27 |
| 39 | Oseltamivir | 3 - 7 dpi, twice a day for the first 3 days and once a day | 60 mg/kg | i.p. | 7 | 3 |
| 40 | Oseltamivir | 4 - 8 dpi, twice a day for the first 3 days and once a day | 60 mg/kg | i.p. | 7 | 4 |
| 41 | Oseltamivir | 2 - 8 dpi, twice a day for the first 3 days, then once a day | 60 mg/kg | i.p. | 15 (2) | 15 |
| 42 | Oseltamivir | 2 - 8 dpi, twice a day for the first 3 days, then once a day | 20 mg/kg | i.p. | 7 | 5 |
| 43 | Oseltamivir | 4 - 6 dpi, once a day | 20 mg/kg | i.p. | 46 (6) | 18 |
| 44 | Oseltamivir | 5 - 7 dpi, once a day | 20 mg/kg | i.p. | 8 | 2 |
| 45 | PAMAM dendrimer | 4 and 5 dpi | 20 mg/kg | i.p. | 10 | 0 |
| 46 | PMX205 | 4 and 5 dpi | 1 mg/kg | Sc. | 10 | 0 |
| 47 | PMX205 | 2 – 7 dpi, once a day | 1 mg/kg | Sc. | 10 | 0 |
| 48 | Ruxolitinib | 4 and 5 dpi, twice a day | 125 mg/kg | Oral | 10 | 0 |
| 49 | Silvestat | 4, 5, and 7 dpi, once a day | 100 mg/kg | i.p. | 10 | 0 |
| 50 | Silvestat | 2 – 7 dpi, once a day | 100 mg/kg | i.p. | 10 | 0 |
| 51 | Topotecan Hydrochloride | 4 and 5 dpi, once a day | 2 mg/kg | i.p. | 10 | 0 |
| 52 | Ziluton | 2 – 7 dpi, once a day | 60 mg/kg | Sc. | 10 | 1 |
| 53 | Anti-CD40 | 2 dpi, once a day | 100 µg/mouse | i.p. | 10 | 0 |
|  | Anti-IL-6 | 3 – 8 dpi, once a day | 200 µg/mouse | i.p. |  |  |
| 54 | Anti-IL-6 | 2 – 4 dpi, once a day | 100 µg/mouse | i.p. | 10 | 0 |
|  | Silvestat | 5 – 8 dpi, once a day | 100 mg/kg | i.p. |  |  |
| 55 | Anti-Ly6G | 1 dpi, once a day | 100 µg/mouse | i.p. | 10 | 0 |
|  | Baricitinib | 5 and 7 dpi | 50 mg/kg | Sc. |  |  |
| 56 | AZD-5069 | 1 – 3 dpi, once a day | 100 mg/kg | i.p. | 10 | 0 |
|  | Anti-PSLG-1 |  | 200 µg/mouse |  |  |  |
| 57 | Baricitinib | 2 – 7 dpi, once a day | 25 mg/kg | Sc. | 10 | 0 |
|  | Anti-IL-6 | 2 – 7 dpi, once a day | 200 µg/mouse | i.p. |  |  |

**Table S3.** Reagent list for flow cytometry and histology.

| Antibody name | Make | Clone | Catalog number | RRID |
| --- | --- | --- | --- | --- |
| Anti-mouse CD45 | BioLegend | 30-F11 | 103116 | AB_312981 |
| Anti-mouse CD11b | BioLegend | M1/70 | 101235 | AB_10897942 |
| Anti-mouse CD11c | BioLegend | N418 | 117311 | AB_389306 |
| Anti-mouse NK1.1 | BioLegend | PK136 | 108716 | AB_493590 |
| Anti-mouse Siglec-F | BD Bioscience | E50-2440 | 562680 | AB_2687570 |
| Anti-mouse Ly6G | BD Bioscience | 1A8 | 563978 | AB_2716852 |
| Anti-mouse MHCII | BioLegend | M5/114.15.2 | 107641 | AB_2565975 |
| Anti-mouse CD3 | BioLegend | 17A2 | 100206 | AB_312663 |
| Anti-mouse CD4 | BioLegend | GK1.5 | 100434 | AB_893324 |
| Anti-mouse CD8 | BioLegend | 53-6.7 | 100722 | AB_312761 |
| Anti-mouse CD19 | BD Bioscience | 1D3 | 562291 | AB_11154223 |
| Anti-mouse CD16/32 | BioLegend | 93 | 101302 | AB_312801 |
| Anti-mouse Ly6C | BioLegend | HK1.4 | 128018 | AB_1732082 |
| Zombie UV Fixable Viability Kit | BioLegend |  | 423108 |  |
| Anti-Influenza A NP | Thermo Fisher Scientific | D67J | MA1-7322 | AB_1017747 |
| Hoechst 333532 | Biotium |  | 40044 |  |
| Anti-Pro surfactant protein C | Abcam | Rabbit polyclonal | ab90716 |  |
| Anti-mouse EpCAM | BioLegend | G8.8 | 118222 | AB_2563322 |
| Anti-mouse EpCAM | BioLegend | G8.8 | 118240 | AB_2810354 |
| Anti-mouse Ly6G | BioLegend | 1A8 | 127626 | AB_2561340 |
| Anti-mouse Ly6G | BioLegend | 1A8 | 127610 | AB_1134159 |
| Anti-mouse CD11b | BioLegend | M1/70 | 101254 | AB_2563231 |
| Anti-Alpha smooth muscle actin | Thermo Fisher Scientific | 1A4 | 53-9760-82 | AB_2574461 |
| Anti-mouse Arginase-1 | Thermo Fisher Scientific | A1exF5 | 12-3697-82 | AB_2734839 |
| Anti-Ki67 | BD Bioscience | B56 | 561126 | AB_10611874 |
| Anti-Phospho-Stat3 (Tyr705) | Cell Signaling Technology | D3A7 | 9145S |  |
| Mouse Fc Block | BD Bioscience | 2.4G2 | 553142 | AB_394656 |
| Donkey anti-Rabbit IgG | Thermo Fisher Scientific | Donkey polyclonal | A-31572 | AB_162543 |
| Goat Anti-Rabbit IgG | Abcam | Goat polyclonal | ab175735 |  |

**Movie S1.** Clearing-enhanced 3D (Ce3D) imaging of the lung infected with PR8 at 3 DPI. Cyan, green, magenta, and white represent EpCAM, IAV nucleoprotein (NP), Ly6G, and CD11b, respectively.

